# Multi-Omic Profiling Defines the Renal Mechanisms of Cardiovascular-Kidney-Metabolic Syndrome in Pulmonary Arterial Hypertension

**DOI:** 10.64898/2026.09.23.753744

**Authors:** Madelyn J. Blake, Sally E. Prins, Jenna B. Mendelson, Lynn M. Hartweck, Todd Markowski, LeeAnn Higgins, Sandra Breulis-Bonnet, Steeve Provencher, Sébastien Bonnet, Kurt W. Prins

## Abstract

**Background:** Cardiovascular-kidney-metabolic (CKM) syndrome integrates cardiac, renal, and metabolic abnormalities that drive multi-organ injury. In pulmonary arterial hypertension (PAH), CKM manifests as systemic metabolic derangements, right ventricular dysfunction, and renal compromise. Renal dysfunction strongly predicts mortality in PAH, yet current therapies provide little renal benefit. Moreover, the mechanisms driving PAH nephropathy remain poorly understood, limiting our ability to effectively treat PAH-CKM.

**Methods:** Single-nucleus RNAsequencing identified cell-type-specific transcriptional alterations in autopsy-derived kidneys from control (n=5) and PAH (n=4) patients. Mitochondrial, cytoplasmic, and phosphoproteomic analyses profiled proteomic alterations. AlphaFold 3/ChimeraX modeling defined the predicted structural consequences of altered protein phosphorylation. Histological analysis assessed renal fibrosis, glomerular structure, immune cell infiltration, and nephron segment density.

**Results:** 3 of 4 PAH patients exhibited renal impairment, with a cohort mean estimated glomerular filtration rate of 61±36 mL/min/1.73m². snRNA-seq identified a distinct cellular architecture in PAH kidneys, marked by an increase in thick ascending limb, proximal tubule, and immune cell nuclei and depletion of collecting duct nuclei. Transcriptional profiling demonstrated proximal tubule and thick ascending limb cells both upregulated fatty acid oxidation, ferroptosis, and cuproptosis pathways. PAH lymphocytes displayed heightened T-cell receptor signaling, natural killer cell-mediated cytotoxicity, and Th17 differentiation. Histological analyses demonstrated increased perivascular fibrosis, glomerular T-cell infiltration, and a reduction in glomerular basement membrane density in PAH kidneys. Mitochondrial and cytoplasmic proteomics revealed broad metabolic dysfunction characterized by impaired β-oxidation, TCA cycle activity, amino acid metabolism, transsulfuration, and urea cycle pathways. Phosphoproteomics predicted increased GSK3β, STK, and casein kinase activity in PAH kidneys. Finally, using the totality of our data, we nominated multiple druggable targets that could be evaluated to counteract PAH nephropathy.

**Conclusions:** Integrated multi-omic profiling demonstrates PAH nephropathy is defined by glomerular structural remodeling, proximal nephron ferroptotic and cuproptotic signaling, amino acid metabolic dysregulation, and innate and adaptive leukocyte activation. Future studies intervening on these pathways could lead to the development of novel therapeutics to augment renal function in PAH.

**Clinical Perspective:** *What Is New?:*

- Single-nucleus RNA sequencing and histological analyses defined nephron remodeling and increased leukocyte infiltration as key cellular features of the PAH kidney.
- Integrated transcriptomic and proteomic analyses nominated metabolic derangements including ferroptosis, cuproptosis, disrupted fatty acid and amino acid metabolism, and impaired TCA cycle activity as candidate drivers of PAH nephropathy.

*What Are the Clinical Implications?:*

- Discrete cellular and metabolic mechanisms may contribute to PAH-CKM syndrome, which suggests renal dysfunction is not solely due to abnormal hemodynamics.
- Dysregulated programmed cell death pathways, immune signaling, amino acid metabolism, and kinase activity may be potentially druggable targets that could be engaged to counteract PAH-CKM.

## Introduction

Cardiovascular-kidney-metabolic (CKM) syndrome is a systemic disorder linking cardiac, renal, and metabolic abnormalities that ultimately drive multi-organ injury and mortality^1^. Features of CKM syndrome frequently coexist in PAH, including systemic metabolic abnormalities, RV dysfunction, and renal impairment^2^. Clinical markers of kidney injury, including elevated serum creatinine and albuminuria, are associated with RV dysfunction, systemic inflammation, insulin resistance, and increased mortality in PAH^3,4^. Notably, an analysis of 6,694 patients across 18 phase III pulmonary hypertension trials demonstrated that impaired renal function was associated with a 16% increase in all-cause mortality^5^. Unfortunately, current PAH therapies minimally improve renal function^5^. These clinical data underscore the pressing need to delineate the cellular and molecular mechanisms underlying renal dysfunction in PAH so novel therapeutic approaches can be pursued.

Lymphocytes contribute to renal injury across multiple forms of nephropathy. In murine hypertension-associated chronic kidney disease, inhibition of cytokine-producing T-cell infiltration into the tubulointerstitum prevents glomerular injury and fibrosis^6^. Furthermore, IL-17-producing lymphocytes have also been implicated in renal fibrosis following ischemia-reperfusion injury^7^ and obstructive nephropathy^8^. In addition to T cells, natural killer (NK) cells induce tubular epithelial apoptosis in experimental kidney injury^9^. In agreement with these findings, activated NK cell abundances correlate with renal dysfunction in human fibrotic kidney disease^10^. Collectively, these preclinical and clinical studies implicate lymphocytes as key mediators of kidney injury; however, their role in PAH nephropathy remains unknown.

Metabolic stress-induced death pathways, particularly ferroptosis and cuproptosis-related signaling are implicated in kidney disease across diverse renal insults. In ischemia-reperfusion, ferroptosis drives proximal tubular loss, while its pharmacologic inhibition preserves renal function^11^. Cuproptosis is both a biomarker and pathogenic mediator of renal disease, with associated gene signatures predicting outcomes in clear cell renal cell carcinoma and diabetic kidney disease^12^. Moreover, excess intracellular copper promotes fibrotic remodeling through enhanced lysyl oxidase activity in preclinical models of kidney fibrosis^13^. Finally, cuproptosis-associated gene signatures correlate with immune-cell infiltration and markers of disease severity in diabetic kidney disease^14^. Collectively, these findings implicate ferroptosis and cuproptosis as key drivers of renal injury, inflammation, and fibrosis; however, whether these pathways contribute to PAH nephropathy is unclear.

Here, we aimed to define the renal mechanisms underlying CKM syndrome in PAH through integrated multi-omic and histological analyses of autopsy-derived kidneys from PAH patients and non-PAH controls. Single-nucleus RNA sequencing (snRNA-seq) characterized the cellular landscape and cell-type-specific transcriptional alterations, while three proteomic analyses identified pathways dysregulated with renal dysfunction. Finally, we leveraged these multi-omic findings to nominate readily translatable therapeutic strategies for PAH-CKM syndrome.

## Methods

### Human Tissue Collection and Study Approval

Autopsy-derived kidney tissues were obtained from 5 non-PAH control donors and 4 patients with PAH at Laval University and stored at −80°C until analysis. Clinical characteristics are provided in **Supplemental Table 1**. Tissue use conformed to the Declaration of Helsinki and was approved by the Centre de recherche de l’Institut de cardiologie et de pneumologie de Québec ethics committee (CER 20773). Written informed consent was obtained from patients before death or, when this was not possible, from their next of kin.

### Single-Nucleus RNA Sequencing

Nuclei were isolated from frozen kidney tissues, purified by flow cytometry, and processed using the 10x Genomics platform^15^. Libraries were sequenced and aligned to the GRCh38 human reference genome. Gene-expression matrices were processed in Seurat^15^ using sample-specific quality-control thresholds, normalization, principal component analysis, DoubletFinder^16,17^-based doublet removal, and graph-based clustering. Cell identities were assigned using Azimuth mapping to a human kidney reference^18^ and confirmed using conserved renal markers and the Human Protein Atlas^19^. Differential expression was performed on sample-pseudobulked counts using DESeq2; genes with an absolute log2 fold change ≥0.50 and a Benjamini–Hochberg-adjusted P<0.05 were considered differentially expressed. Pathway enrichment was assessed using ShinyGO^20^ with KEGG and WikiPathways annotations, and pathway module scores were calculated from sample-aggregated expression within individual cell populations. MHC class I machinery and classical HLA-A, HLA-B, and HLA-C expression were evaluated by cell type. Intercalated-to-principal-cell trajectories and Notch-associated transcriptional programs were examined using Monocle 3^21^.

### Kidney Histology and Immunofluorescence

Paraffin-embedded kidney sections were stained with Masson’s trichrome, and total and perivascular fibrosis were quantified blinded using ImageJ. Confocal immunofluorescence identified proximal tubules with Lotus tetragonolobus lectin, thick ascending limbs with uromodulin, collecting ducts with Dolichos biflorus agglutinin, T cells with CD3, and NK cells with CD57. Images were acquired under identical conditions, and measurements were performed blinded in FIJI. Replicate measurements were summarized to provide one value per donor, except where individual regions are specified in the figure legends.

### Proteomic and Phosphoproteomic Analyses

Mitochondria-enriched^22^ and cytoplasmic^23^ kidney fractions underwent tandem mass tag labeling and quantitative mass spectrometry. Relative pathway abundance was calculated by summing detected protein abundances assigned to predefined KEGG or WikiPathways pathways and normalizing values to the control mean. Phosphopeptide-enriched kidney samples were analyzed by mass spectrometry, and phosphoproteins differing between PAH and control kidneys at adjusted P<0.05 were evaluated using Kinase Enrichment Analysis 3^24^. The highest-ranked predicted kinases were visualized using Coral^25^.

### Protein Structural Modeling

Native and phosphorylated forms of 41 differentially phosphorylated proteins were modeled using AlphaFold 3^26^ (version 3.0.1) and analyzed in UCSF ChimeraX^27^ (version 1.8). Models were prioritized according to phosphosite confidence, proximity to functionally characterized residues, and biological relevance. Native and phosphorylated structures were compared using structural alignment, root-mean-square deviation, electrostatic surface potential, solvent-accessible surface area, hydrogen-bond, and atomic-clash analyses. Because HAO2 pS349 lies within its C-terminal peroxisomal targeting signal, focused models of the native or phosphorylated HAO2 C-terminal peptide bound to the PTS1-recognition domain of PEX5 were used for quantitative interface analyses. Full-length HAO2–PEX5 models provided structural context. Complete modeling and model-selection procedures are described in the Supplemental Methods.

### Statistical Analysis

The biological donor was used as the unit of analysis unless otherwise indicated. Normality was evaluated using the Shapiro–Wilk test, followed by an unpaired Student’s t test or Mann–Whitney U test, as appropriate. Differential-expression P values were corrected using the Benjamini–Hochberg method. Tests were 2-sided, and P<0.05 was considered statistically significant. Data are presented as means or medians, as appropriate, with individual values shown. Analyses and graphing were performed using GraphPad Prism version 10 and R.

### Data Availability

Raw snRNA-seq data are deposited in the NCBI Gene Expression Omnibus, proteomic and phosphoproteomic data are available through Zenodo (DOI:10.5281/zenodo.21629606), and analysis code is available on GitHub (https://github.com/blake561/Multi-Omic-Profiling-Defines-the-Renal-Mechanisms-of-Cardiovascular-Kidney-Metabolic-Syndrome-in-PAH). Additional materials are available upon reasonable request.

## Results

### Histological Evaluation Identified Heightened Perivascular Fibrosis, Impaired Glomerular Structure, and Altered Nephron-Segment Abundances

We first characterized the histopathologic features of PAH-associated kidney disease. Masson’s trichrome staining revealed no difference in overall renal fibrosis between groups (**Figure 1A**); however, perivascular fibrosis was increased in PAH kidneys (**Figure 1B**). Gross histological assessment showed no change in glomerular density (**Figure 1C**), whereas high-resolution imaging demonstrated reduced glomerular basement membrane density (**Figure 1D**). We next evaluated the abundance of proximal convoluted tubules, thick ascending limbs, and collecting ducts by immunohistochemistry. Although these differences did not reach statistical significance, PAH kidneys had a nonsignificant increase in proximal convoluted tubule and thick ascending limb abundances, and a nonsignificant reduction in collecting duct density (**Figure 1E–G**). Collectively, these findings demonstrated PAH kidneys exhibited localized fibrotic remodeling, glomerular structural alterations, and a potential loss of collecting ducts.

**Figure 1.**
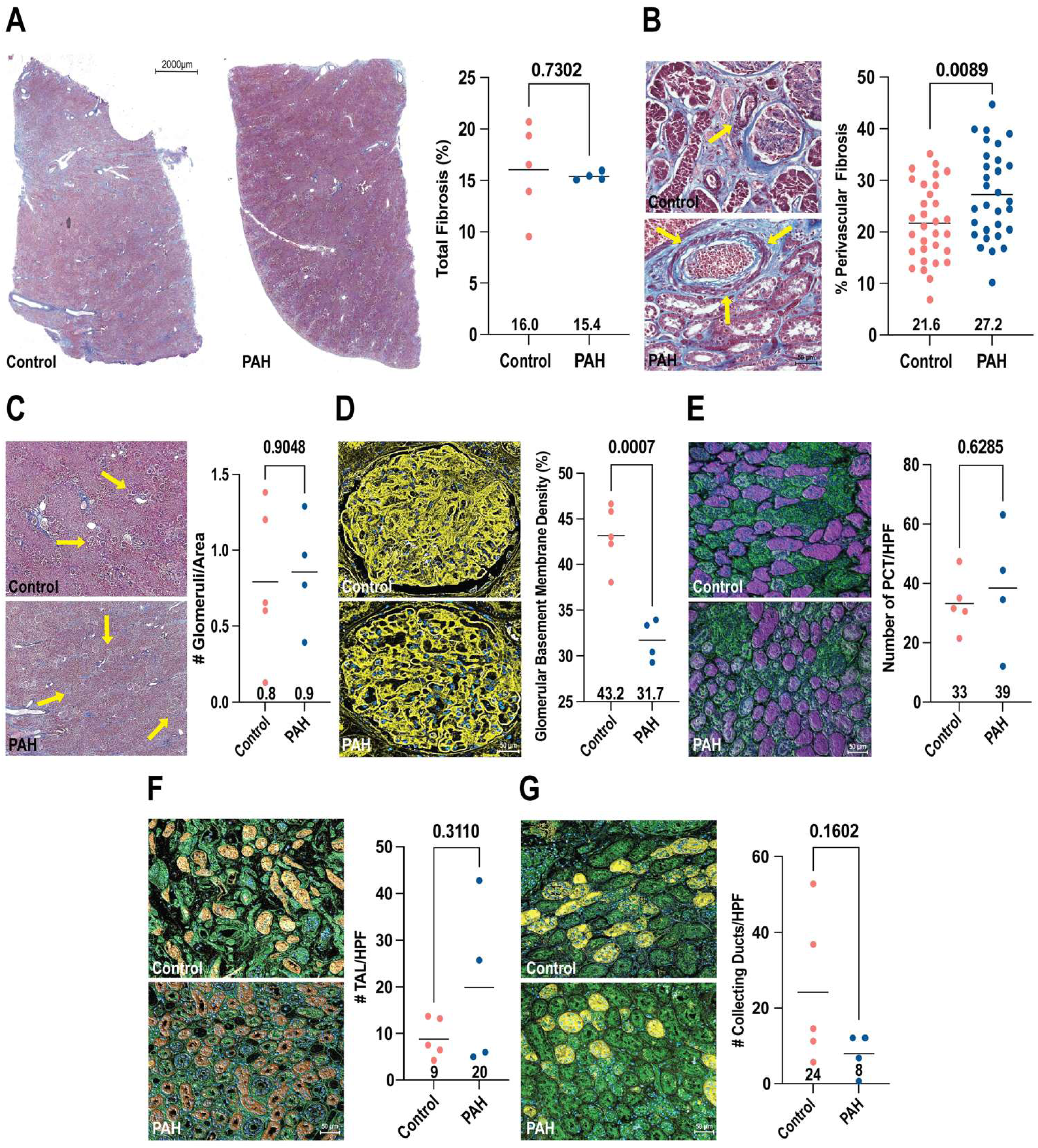
PAH kidneys exhibit increased perivascular fibrosis, reduced glomerular basement membrane density, and remodeling of nephron-segment abundance. (A) Representative whole-section Masson’s trichrome images and blinded quantification of total renal fibrosis in control and PAH kidneys. (B) Representative Masson’s trichrome images and quantification of perivascular fibrosis. Yellow arrows indicate perivascular collagen. (C) Representative renal sections and quantification of glomerular density. Yellow arrows indicate glomeruli. (D) Representative confocal images and quantification of glomerular basement membrane density. (E) Representative confocal images and numbers per high-power field (HPF) of lotus tetragonolobus lectin–positive proximal convoluted tubules (PCTs), (F) uromodulin-positive thick ascending limbs (TALs), and (G) Dolichos biflorus agglutinin– positive collecting ducts. Control, n = 5; PAH, n = 4. In B, each point represents an analyzed perivascular region; in A and C–G, each point represents one patient. Horizontal lines indicate medians. *P* values were calculated using two-sided Mann–Whitney U tests and are shown above each comparison.

### Single-Nucleus RNA Sequencing Defined the Cellular Architecture and Transcriptomic Alterations of the PAH Kidney

To define the cellular and molecular changes potentially underlying these histological findings, we performed single-nucleus RNA sequencing of PAH and control kidneys. Analysis of 32,331 control and 56,993 PAH nuclei identified 11 distinct renal cell types (**Supplemental Figure 2A and B**). PAH kidneys exhibited greater relative abundances of thick ascending limb (14.2% vs. 43.4%), proximal tubule (7.0% vs. 14.3%), parietal cell (1.0% vs. 2.7%), lymphocyte (0.8% vs. 2.4%), and macrophage (0.3% vs. 0.8%) populations. Conversely, the relative abundances of principal cell (38.2% vs. 17.1%), intercalated cell (8.4% vs. 1.3%), distal convoluted tubule (10.0% vs. 5.0%), podocyte (2.0% vs. 0.1%), endothelial cell (12.7% vs. 8.2%), and fibroblast (5.5% vs. 4.7%) populations were reduced (**Supplemental Figure 2C**). Principal component analysis of sample-level transcriptomes further demonstrated separation between control and PAH kidneys (**Supplemental Figure 2D**). Differential gene expression analysis revealed extensive transcriptional alterations, most prominently in proximal tubule, thick ascending limb, and principal cells (**Supplemental Figure 2E**). Among non-nephron populations, lymphocytes and endothelial cells exhibited the most substantial transcriptional alterations, whereas macrophage and podocyte transcriptomes remained comparatively stable (**Supplemental Figure 2E**).

### PAH Manifested as a Pro-Ferroptotic/Cuproptotic Transcriptional State in the Proximal Convoluted Tubule and Thick Ascending Limb

Because proximal tubule and thick ascending limb populations were expanded and extensively transcriptionally altered in PAH kidneys, we examined these cell types in greater depth (**Figure 2A**). UMAP projections of proximal tubule cells demonstrated segregation between control and PAH samples (**Figure 2B**). PAH kidneys exhibited enrichment of fatty acid oxidation, ferroptosis, and copper homeostasis pathways on pathway analysis and this was accompanied by increases in fatty acid metabolism, ferroptosis, and cuproptosis module scores (**Figure 2D–E**). Pathways reduced in PAH were heterogeneous across databases and cell types (**Supplemental Figure 3**). In contrast, the thick ascending limb cells showed minimal alteration on UMAP projections (**Figure 2C**), but displayed metabolic remodeling characterized by enhanced oxidative phosphorylation, fatty acid metabolism, and cuproptosis (**Figure 2F–G**).

**Figure 2.**
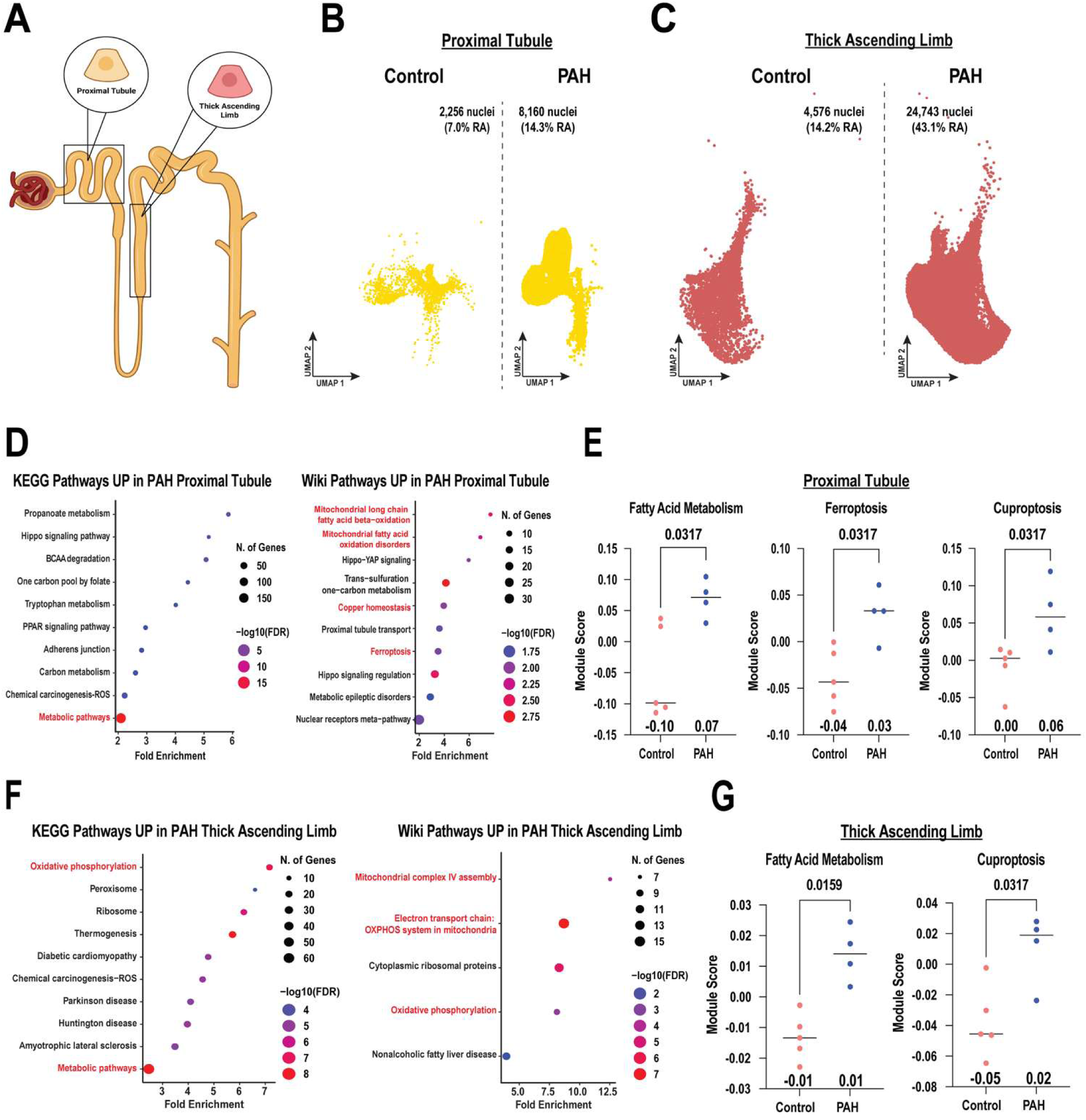
Proximal tubule and thick ascending limb cells in PAH kidneys exhibit metabolic remodeling and increased ferroptosis– and cuproptosis-related programs. (A) Nephron schematic highlighting the proximal tubule and thick ascending limb. (B) UMAP projections of proximal tubule nuclei and (C) thick ascending limb nuclei from control and PAH kidneys; group-specific nuclei counts and relative abundances are indicated. (D) KEGG and WikiPathways pathways enriched in PAH proximal tubule cells relative to controls. (E) Sample-pseudobulked fatty acid metabolism, ferroptosis, and cuproptosis module scores in proximal tubule cells. (F) KEGG and WikiPathways terms enriched in PAH thick ascending limb cells. (G) Sample-pseudobulked fatty acid metabolism and cuproptosis module scores in thick ascending limb cells. In D and F, dot size represents the number of genes and color represents −log10(false-discovery rate [FDR]); selected pathways are highlighted in red. In E and G, each point represents one patient (control, n = 5; PAH, n = 4), horizontal lines indicate group medians, and P values were calculated using two-sided Mann–Whitney U tests. BCAA, branched-chain amino acid; OXPHOS, oxidative phosphorylation; UMAP, uniform manifold approximation and projection.

### PAH Nephropathy Was Characterized by T-Cell and NK-Cell Autoimmune-Like Phenotypes

To investigate immune alterations, we analyzed lymphocyte transcriptomes from control and PAH kidneys. PAH lymphocytes exhibited enhanced T-cell receptor signaling, NK-cell-mediated cytotoxicity, and Th17 differentiation programs, consistent with a pro-inflammatory, cytotoxic phenotype (**Figure 3A–D**). Downregulated pathways were heterogeneous and primarily involved calcium, cytoskeletal, and focal-adhesion signaling (**Supplemental Figure 4**). In agreement with these data, there were increases in CD4/CD8 T cells and NK cells in PAH samples as compared to control (**Supplemental Figure 5**). Given the opposing dependence of cytotoxic T cells and NK cells for MHC class I (MHC-I) engagement (**Figure 3F–G**), we examined MHC-I across renal cell populations. MHC-I expression was reduced in proximal tubule, principal, thick ascending limb, and distal convoluted tubule cells, potentially increasing susceptibility to NK-cell-mediated killing. However, endothelial cells, podocytes, and intercalated cells exhibited elevated levels of MHC-I, which may facilitate CD8 T-cell-mediated death (**Figure 3H**). Supporting these findings, the canonical MHC-I genes *HLA-A*, *HLA-B*, and *HLA-C* were coordinately dysregulated across multiple renal cell populations (**Figure 3I**). Finally, we used confocal microscopy to validate our snRNA-seq findings. PAH kidneys demonstrated significantly greater T-cell infiltration within glomeruli (**Figure 3J**) and there was a nonsignificant trend toward increased NK-cell abundance in the tubulointerstitum (**Figure 3K**).

**Figure 3.**
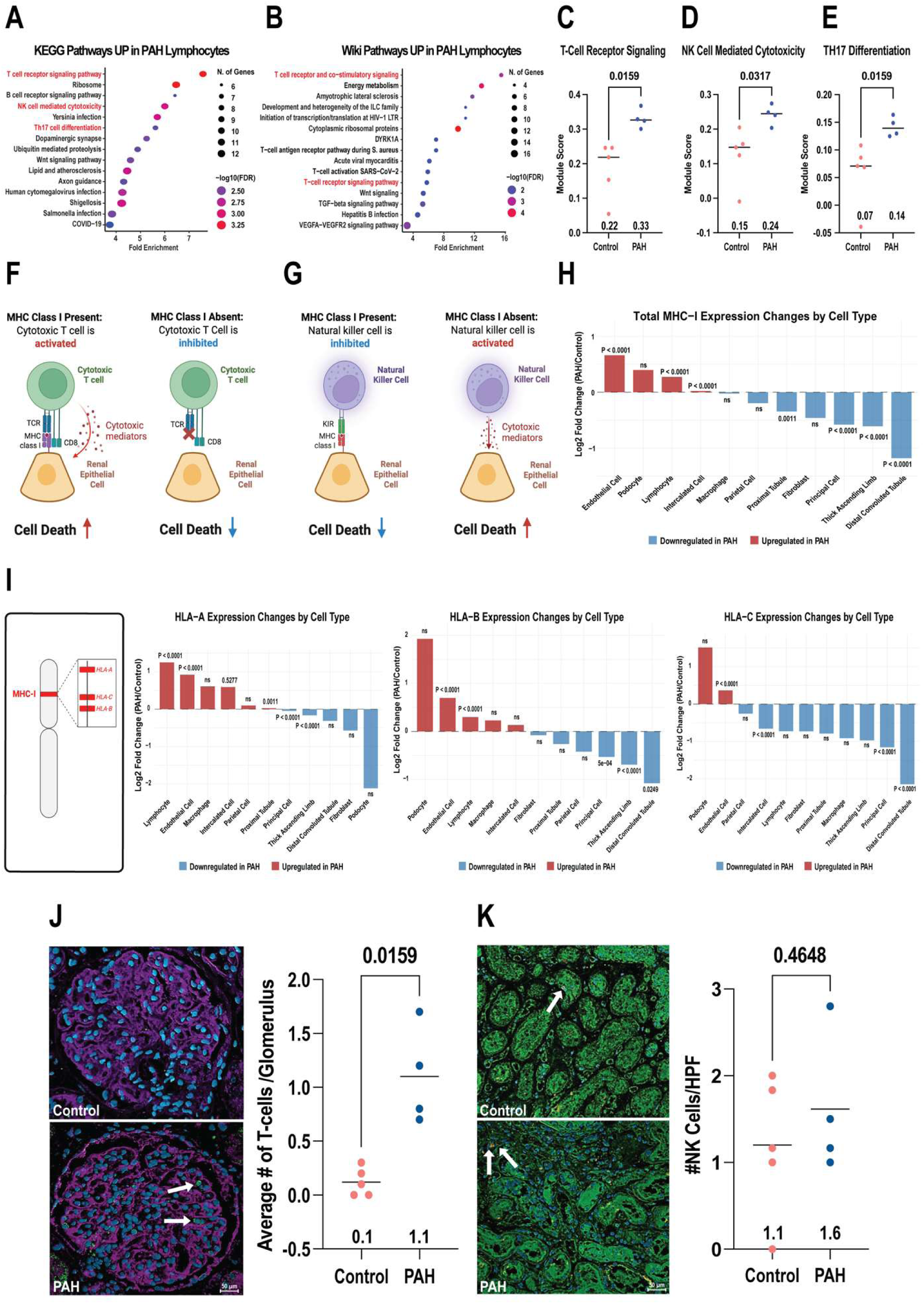
PAH kidneys display lymphocyte activation, cell-type-specific MHC-I dysregulation, and increased glomerular T-cell infiltration. (A) KEGG and (B) WikiPathways terms enriched among genes increased in PAH kidney lymphocytes. Dot size represents the number of genes and color represents −log10(false-discovery rate [FDR]); selected immune pathways are highlighted in red. (C) Sample-pseudobulked module scores for T-cell receptor signaling, (D) natural killer (NK) cell–mediated cytotoxicity, and (E) T helper 17 (Th17) differentiation. Each point represents one patient (control, n = 5; PAH, n = 4), horizontal lines indicate group medians, and P values were calculated using two-sided Mann–Whitney U tests. (F) Schematics illustrating the opposing effects of major histocompatibility complex class I (MHC-I) expression on cytotoxic T-cell activation and (G) NK-cell activation. (H) Log2 fold change in total MHC-I machinery expression in PAH versus control across renal cell types. (I) Chromosomal location of classical MHC-I genes and cell-type-specific log2 fold changes in HLA-A, HLA-B, and HLA-C expression. In H and I, red indicates increased expression and blue indicates decreased expression in PAH; expression was compared using the Wilcoxon rank-sum test, and P values are shown above the bars. (J) Representative confocal images and quantification of CD3-positive T cells per glomerulus. White arrows indicate CD3-positive cells. (K) Representative confocal images and quantification of CD57-positive cells per HPF, used as an estimate of NK-cell abundance. White arrows indicate CD57-positive cells. In J and K, each point represents one patient, horizontal lines indicate medians, and P values were calculated using Mann–Whitney U tests.

### Mitochondria-Enriched Proteomics Identified Potential Metabolic Dysregulation in PAH Kidneys

To supplement our transcriptomic analysis that nominated metabolic dysregulation in the PAH kidney, we evaluated mitochondrial protein remodeling in PAH kidneys by performing quantitative proteomics of mitochondria-enriched fractions. Hierarchical clustering of the top 500 differentially expressed proteins identified a distinct proteomic signature in PAH relative to controls (**Figure 4A**). Pathway analysis demonstrated activation of HIF-1 signaling and downregulation of several metabolic enzymes, including those involved in the TCA cycle, β-oxidation, and multiple amino acid pathways (**Figure 4B–C**). Broader WikiPathways analysis similarly identified reduced mitochondrial fatty acid oxidation, TCA-cycle, and amino acid metabolism pathways, accompanied by enrichment of glycolytic and VEGFA–VEGFR2 signaling programs (**Supplemental Figure 6**). Consistent with these findings, proteomics pathway scores confirmed increased HIF-1 activity (**Figure 4D**) and reduced oxidative metabolism, branched-chain amino acid, tryptophan, and β-alanine metabolism pathways (**Figure 4E–G**).

**Figure 4.**
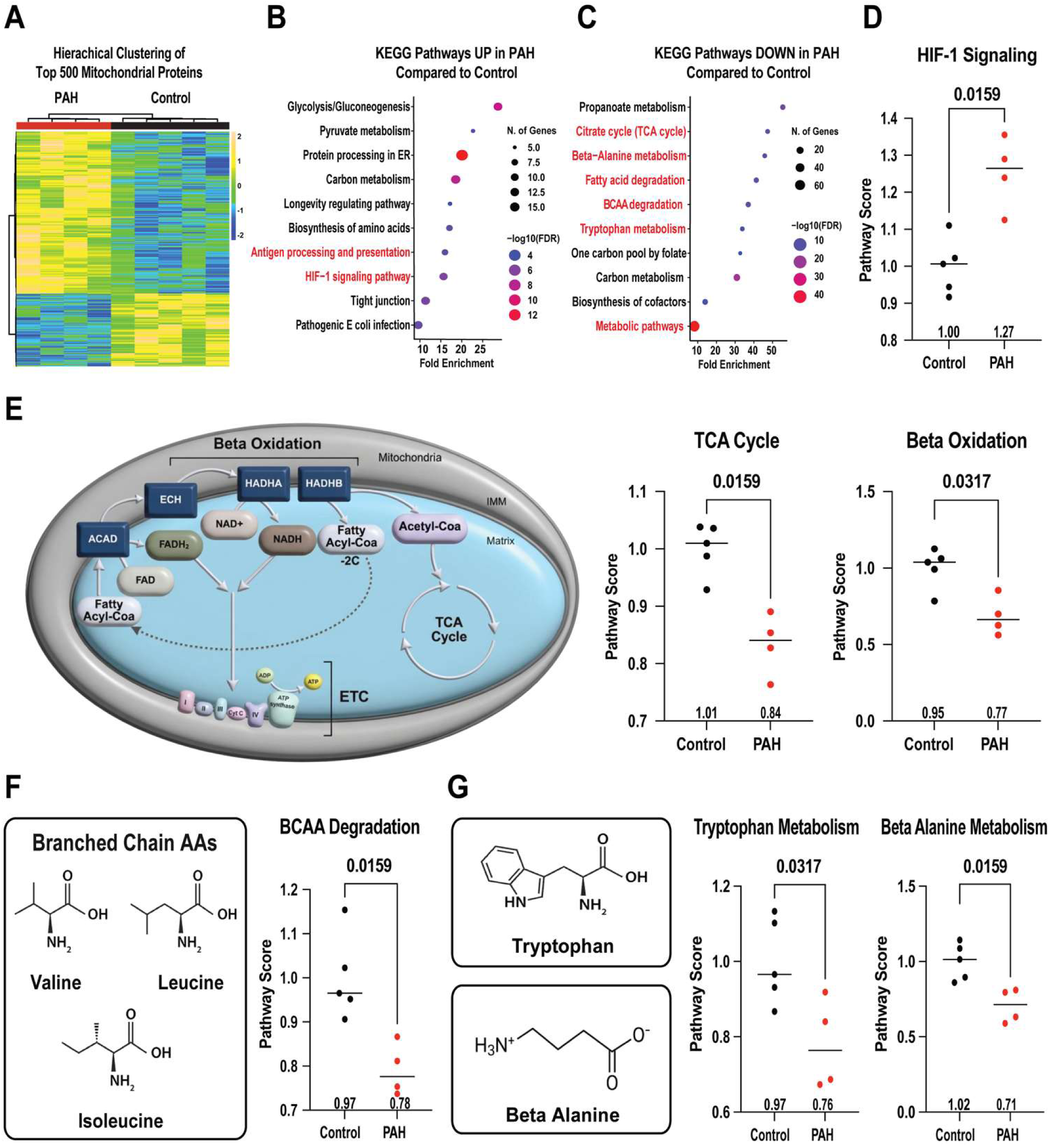
Mitochondria-enriched proteomics identifies HIF-1 activation and impaired oxidative and amino acid metabolism in PAH kidneys. (A) Unsupervised hierarchical clustering of the 500 most variable proteins detected in mitochondria-enriched fractions from control and PAH kidneys. (B) KEGG pathways enriched among proteins increased or (C) decreased in PAH relative to controls. Dot size represents the number of proteins and color represents −log10(false-discovery rate [FDR]); selected pathways are highlighted in red. (D) Relative HIF-1 signaling pathway score. (E) Schematic of mitochondrial β-oxidation, the tricarboxylic acid (TCA) cycle, and the electron transport chain, with relative TCA-cycle and β-oxidation pathway scores. (F) Structures of the branched-chain amino acids valine, leucine, and isoleucine, with the relative branched-chain amino acid degradation score. (G) Structures of tryptophan and β-alanine, with their respective pathway scores. Each point represents one patient (control, n = 5; PAH, n = 4), horizontal lines indicate medians, and P values were calculated using Mann–Whitney U tests. BCAA, branched-chain amino acid; ETC, electron transport chain; HIF-1, hypoxia-inducible factor 1.

### Cytoplasmic Proteomics Delineated Alterations in Amino Acid Homeostasis and Suppressed Transsulfuration

To further increase the depth of our proteomic analysis, we analyzed cytoplasmic fractions from control and PAH kidneys. Hierarchical clustering of the top 500 cytoplasmic proteins identified a distinct proteomic signature in PAH kidneys (**Figure 5A**). Pathway enrichment analysis revealed broad suppression of amino acid metabolism, including the urea cycle, transsulfuration, alanine, aspartate, and glutamate metabolism, and tryptophan metabolism (**Figure 5B–C**). Conversely, proteins increased in PAH were enriched in lipid and cholesterol metabolism and complement-associated inflammatory pathways (**Supplemental Figure 7**). These alterations were confirmed by pathway scores demonstrating reduced cumulative enzyme abundance across each amino acid pathway relative to controls (Figure 5**D–G**).

**Figure 5.**
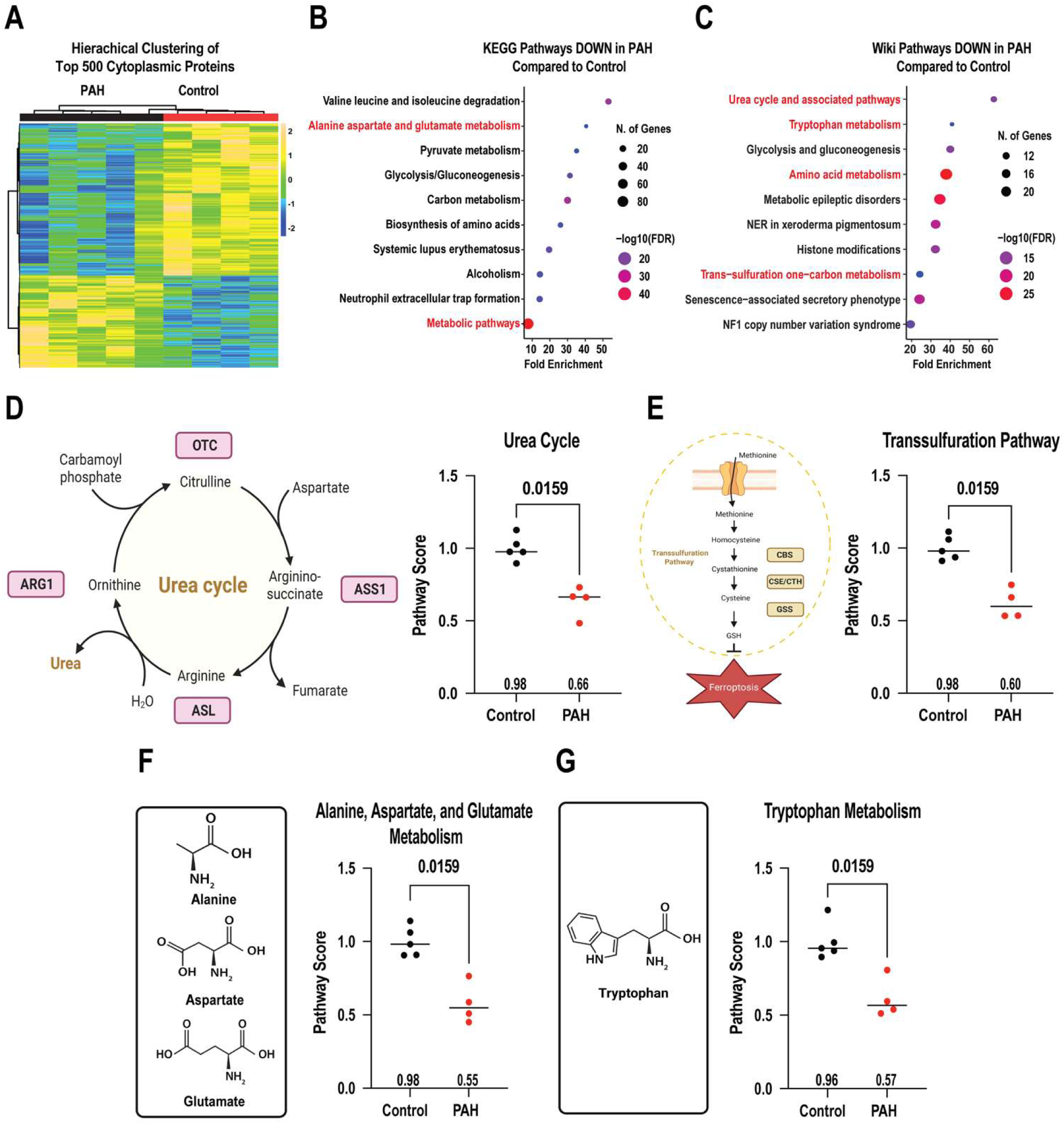
Cytoplasmic proteomics demonstrates broad suppression of amino acid metabolism and transsulfuration in PAH kidneys. (A) Unsupervised hierarchical clustering of the 500 most variable proteins detected in cytoplasmic fractions from control and PAH kidneys. (B) KEGG and (C) WikiPathways pathways enriched among proteins significantly decreased in PAH. Dot size represents the number of proteins and color represents −log10(false-discovery rate [FDR]); selected pathways are highlighted in red. (D) Urea-cycle schematic and relative urea-cycle pathway score. (E) Transsulfuration-pathway schematic and relative transsulfuration pathway score. (F) Structures of alanine, aspartate, and glutamate and the relative alanine, aspartate, and glutamate metabolism pathway score. (G) Tryptophan structure and relative tryptophan metabolism pathway score. Each point represents one patient (control, n = 5; PAH, n = 4), horizontal lines indicate medians, and P values were calculated using Mann–Whitney U tests. ARG1, arginase 1; ASL, argininosuccinate lyase; ASS1, argininosuccinate synthase 1; CBS, cystathionine β-synthase; CSE/CTH, cystathionine γ-lyase; GSH, glutathione; OTC, ornithine transcarbamylase.

### Phosphoproteomics nominated kinase dysregulation and *in silico* simulations predicted structural alterations in metabolic enzymes due to phosphorylation events

Next, we probed how PAH altered post-translational signaling networks by performing quantitative phosphoproteomics. Hierarchical clustering and principal component analysis demonstrated that PAH kidneys exhibited a distinct phosphoproteomic signature compared to controls (**Supplemental Figure 8A–B**). Kinome enrichment analysis predicted activation of the following kinases in PAH kidneys: glycogen synthase kinase 3β (GSK3B), serine/threonine-protein kinases 3 and 4 (STK3 and STK4), SRSF protein kinase 3 (SRPK3), and casein kinase 2 alpha 1 (CSNK2A1) (**Supplemental Figure 8C**). In contrast, kinases predicted to be suppressed were Raf-1 proto-oncogene serine/threonine kinase (RAF1), mitogen-activated protein kinase 8 (MAPK8), AKT serine/threonine kinase 1 (AKT1), glycogen synthase kinase 3α (GSK3A), and mitogen-activated protein kinase 1 (MAPK1) (**Supplemental Figure 8D**).

Then, we performed *in silico* structural analysis of all the differentially phosphorylated proteins in PAH kidneys. Phosphorylated 2-hydroxyacid oxidase 2 (HAO2; pS349), glutathione S-transferase alpha 2 (GSTA2; pS37), and isocitrate dehydrogenase 1 (IDH1; pS94) were predicted to remodel functionally important structural regions. In the HAO2–PEX5 models, the HAO2 C-terminal PTS1 motif localized within the PEX5 binding pocket in both the native and pS349 states. Phosphorylation reorganized the predicted intermolecular hydrogen-bond network, reducing the number of HAO2–PEX5 hydrogen bonds from nine to eight and introducing a phosphate-specific interaction with PEX5 Asn568 (**Supplemental Figure 8E**, **Supplemental Table 3**). GSTA2 pS37 decreased the local mean electrostatic potential near the glutathione binding site and the catalytic residue tyrosine 9 (Y9) from +0.29 to −0.89 (Δ: −1.18 kcal/mol·e) (**Supplemental Figure 8F**). In the IDH1 homodimer, pS94 increased solvent-accessible surface area within 10 Å of the modified residue by 13.9% (native: 6,441 Å²; pS94: 7,336 Å²) and displaced isocitrate-coordinating catalytic residue Arg132 by 0.927 Å relative to the native structure (**Supplemental Figure 8G**). All three phosphosites were less abundant in PAH samples than in controls (**Supplemental Table 3**).

## Discussion

Here, we perform multi-omic profiling of autopsy-derived human kidneys to define the cellular and molecular landscape of the renal component of PAH-associated CKM. Single-nucleus RNA sequencing reveals that PAH alters nephron composition, characterized by a relative increase in thick ascending limb (TAL) and proximal tubule (PT) populations with a reduction of collecting duct cells. PT and TAL cells undergo transcriptional metabolic reprogramming marked by induction of fatty acid oxidation, ferroptosis, and cuproptosis pathways. Furthermore, lymphocytes display enhanced T-cell receptor signaling, NK cell-mediated cytotoxicity, and Th17 differentiation, consistent with a potentially heightened immune activation. Multiple nephron populations also exhibit dysregulated MHC class I expression, which may enhance either NK or T-cell-mediated cytotoxicity. Whole-kidney mitochondrial and cytoplasmic proteomics demonstrate HIF-1 activation and broad suppression of multiple metabolic pathways, including reduced TCA cycle, β-oxidation, and amino acid catabolic pathways. Phosphoproteomics identifies dysregulated kinase signaling and altered phosphorylation of GSTA2, IDH1, and HAO2, which participate in oxidative stress defense, TCA-cycle metabolism, and peroxisomal fatty-acid metabolism, respectively. Structural modeling predicts that these phosphorylation changes may remodel functionally important protein regions, including the HAO2–PEX5 targeting interface. Histological and confocal analyses corroborate many of the molecular findings, including an increase in perivascular fibrosis, a reduction in glomerular basement membrane density, loss of collecting ducts, and T-cell and NK-cell infiltration. Collectively, these data nominate metabolic reprogramming of PT and TAL cells that promotes ferroptosis and cuproptosis, immune activation with dysregulated MHC class I homeostasis, and widespread suppression of renal metabolic pathways as mechanistic drivers of PAH nephropathy.

The kidney regulates systemic physiology through functions that extend well beyond maintenance of fluid and electrolyte homeostasis, and thus the kidney may contribute to some of the systemic manifestations of CKM in PAH. In particular, the proximal convoluted tubule serves as a major site of gluconeogenesis and can account for up to 50% of endogenous glucose production during prolonged fasting or other metabolic stressors^28^. Our single-nucleus RNA sequencing data reveal activation of gluconeogenic programs in the PCT, a finding corroborated by whole-kidney proteomics (**Supplemental Figure 6**). These observations may help explain the observation that PAH patients have insulin resistance and abnormalities in glucose homeostasis^29^. Moreover, our previous work demonstrates that hepatic gluconeogenesis is suppressed in PAH^15^, suggesting that enhanced renal gluconeogenesis may represent a compensatory mechanism and the kidney could be the predominant source of excess gluconeogenesis in PAH patients. This response is distinct from chronic kidney disease, as renal gluconeogenesis is suppressed and is believed to contribute to an increased risk of hypoglycemia^30^. Beyond glucose metabolism, the kidney also plays a central role in kynurenine metabolism^31^. Kynurenine, a product of tryptophan catabolism, accumulates in patients with PAH, and higher circulating levels associate with greater disease severity and reduced survival^32^. When probing our proteomics data, kynureninase and 3-hydroxyanthranilate 3,4-dioxygenase, two key enzymes for kynurenine catabolism, are reduced in PAH samples (**Supplemental Figure 9**). Reduced abundance of these enzymes could promote accumulation of toxic kynurenine intermediates, including 3-hydroxykynurenine. Interestingly, 3-hydroxykynurenine is elevated in PAH serum samples^32^, so impaired renal kynurenine metabolism could underlie this observation. Supporting a pathogenic role for the kynurenine pathway in PAH, pharmacologic inhibition of indoleamine 2,3-dioxygenase 1, the enzyme that converts tryptophan to kynurenine, lowers downstream kynurenine metabolites and attenuates pulmonary vascular remodeling in monocrotaline rats^33^. Together, these findings suggest that metabolic dysfunction in the PAH kidney may contribute to systemic disease by altering glucose homeostasis and kynurenine metabolism.

Another mechanism by which renal dysfunction impairs systemic physiology is through altered handling of proteogenic and non-proteogenic amino acids^34^. Under normal conditions, amino acids are freely filtered by the glomerulus, and the proximal tubule reabsorbs approximately 97%–98% of the filtered load^34^. As renal function declines, impaired tubular reabsorption increases urinary amino acid losses, which may then contribute to skeletal muscle atrophy and reduced exercise capacity in PAH. In addition, we found that branched-chain amino acid metabolism was suppressed in PAH kidneys, and this may have relevance to pulmonary vascular remodeling as we recently demonstrated excess branched-chain amino acids can promote pulmonary vascular remodeling^22^. Beyond proteogenic amino acids, our proteomic analysis reveals suppression of β-alanine metabolism in PAH kidneys (**Figure 4G**). β-Alanine combines with histidine to form carnosine, a dipeptide that enhances exercise performance by buffering intracellular pH, limiting oxidative stress, and facilitating calcium release from the sarcoplasmic reticulum in skeletal muscle^35^. In PAH, circulating β-alanine levels are reduced, and β-alanine supplementation mitigates pulmonary vascular remodeling and right ventricular dysfunction in the monocrotaline rat model of PAH^36^. Together, these observations raise the possibility that impaired renal amino acid handling contributes to exercise intolerance and disease progression in PAH. However, future studies are needed to determine whether renal amino acid wasting, and impaired metabolism directly drive these systemic manifestations.

While our snRNA-seq data demonstrate that immune cells and nephron segments are major contributors to PAH nephropathy, we also observed substantial transcriptional alterations in renal endothelial cells and fibroblasts. Similar to other cell populations, endothelial cells exhibited evidence of metabolic remodeling, including enhanced fatty acid metabolism, ferroptosis, and cuproptosis pathways (**Supplemental Figure 10**). Interestingly, endothelial cell ferroptosis perpetuates glomerular disease in a rodent model of antineutrophil cytoplasmic antibody-induced crescentic glomerulonephritis^37^, suggesting endothelial cell ferroptosis may be another mechanism that drives renal dysfunction and glomerular alterations in PAH. These findings further support the emerging concept that endothelial dysfunction in PAH extends beyond the pulmonary vasculature, consistent with observations in the heart, skeletal muscle, and liver^2^. In addition, PAH renal fibroblasts possess a pro-ferroptotic transcriptional signature that is accompanied by heightened inflammatory signaling via the JAK/STAT pathway. Extensive evidence implicates JAK/STAT signaling in the development of renal fibrosis across multiple forms of nephropathy^38^, raising the possibility that this pathway contributes to the increased perivascular fibrosis observed in PAH kidneys.

Finally, by integrating our multi-omic datasets, we identify several potentially druggable pathways for PAH-associated CKM. Importantly, many of these pathways are already targeted by FDA-approved therapies or clinically advanced compounds, creating opportunities for rapid translational testing. Ferroptosis and cuproptosis emerge as prominent therapeutic targets and both pathways can be attenuated using iron and copper chelators, including deferoxamine, tetrathiomolybdate, and trientine, all of which demonstrate renoprotective effects in experimental kidney injury^13,39,40^ (**Table 1**). Immune dysregulation also represents a promising target, and antibodies directed against NKG2D or interleukin-17, as well as low-dose IL-2 therapy to restore regulatory T-cell homeostasis, show efficacy in models of renal inflammatory injury^9,41,42^ (**Table 1**). Likewise, the kynurenine pathway can be modulated with epacadostat, an indoleamine 2,3-dioxygenase 1 inhibitor, or the kynurenine monooxygenase inhibitor RO 61-8048, both of which exhibit renoprotective properties^43,44^. Finally, dysregulated kinase signaling may be amenable to pharmacologic inhibition, as inhibitors of GSK3β^45^ and CK2α^46^ combat renal injury in preclinical studies. Collectively, these findings identify multiple therapeutic strategies that warrant evaluation in PAH-associated CKM syndrome, particularly given that current PAH therapies provide little protection against progressive renal dysfunction^5^.

**Table 1.**
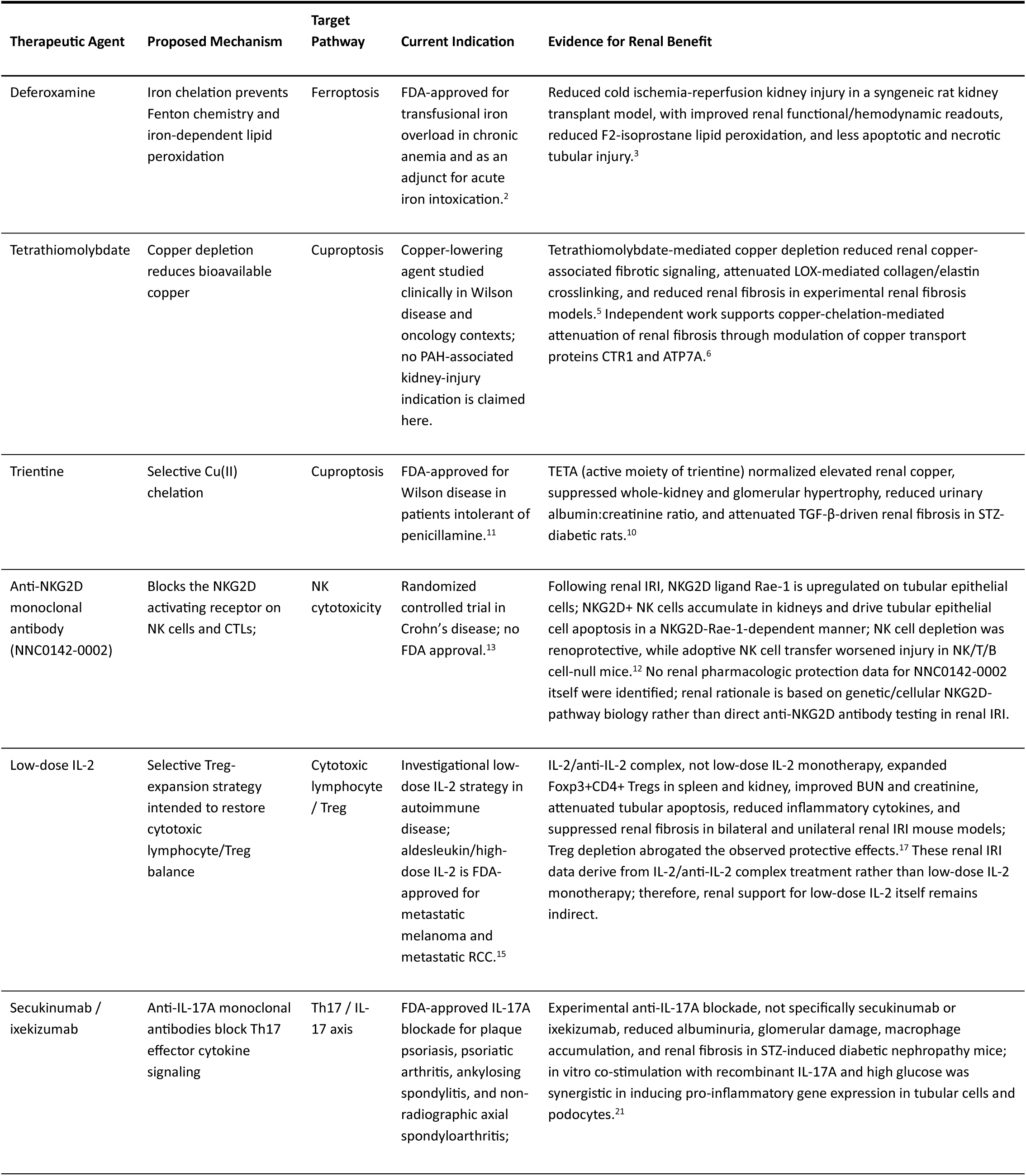
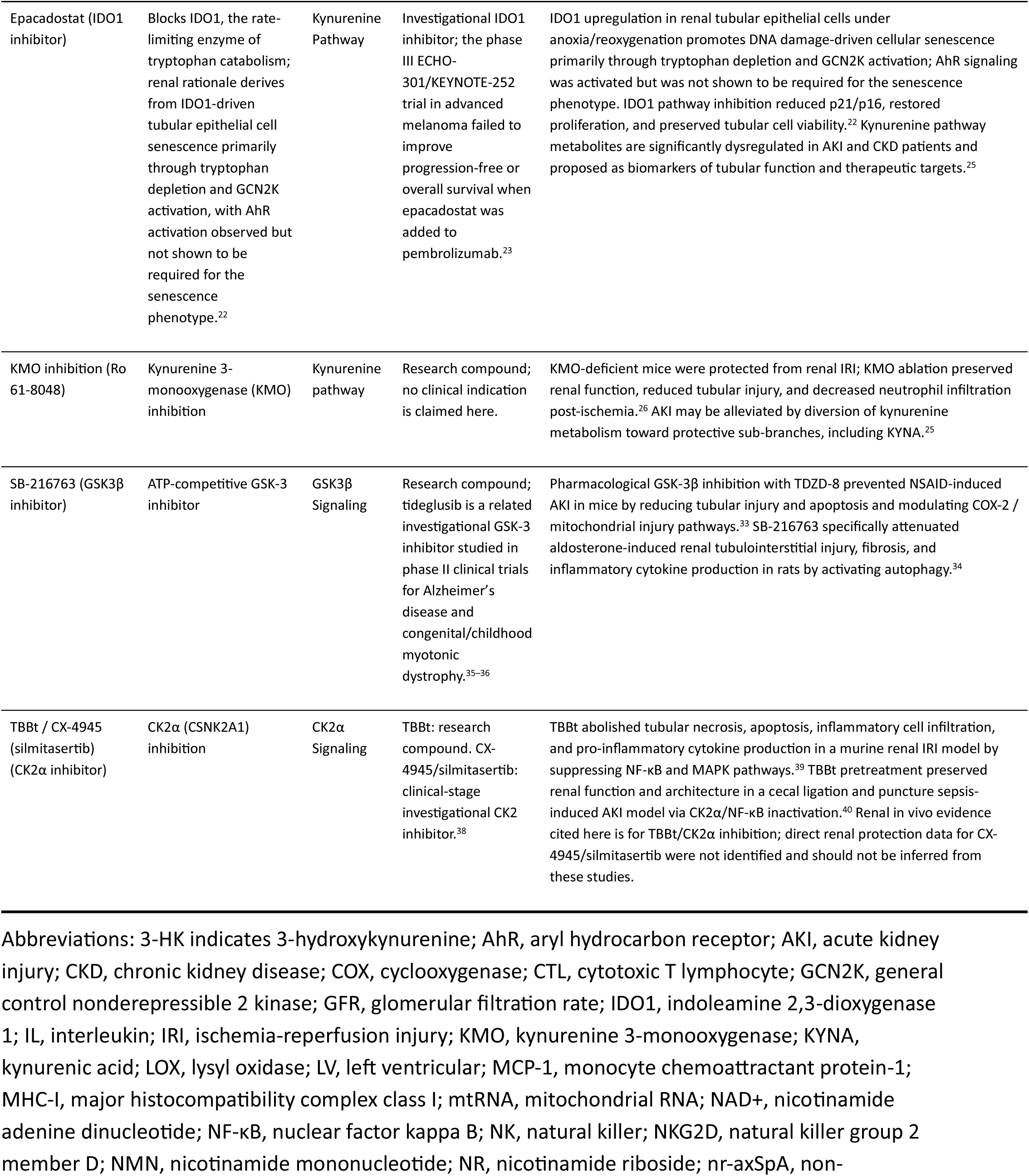
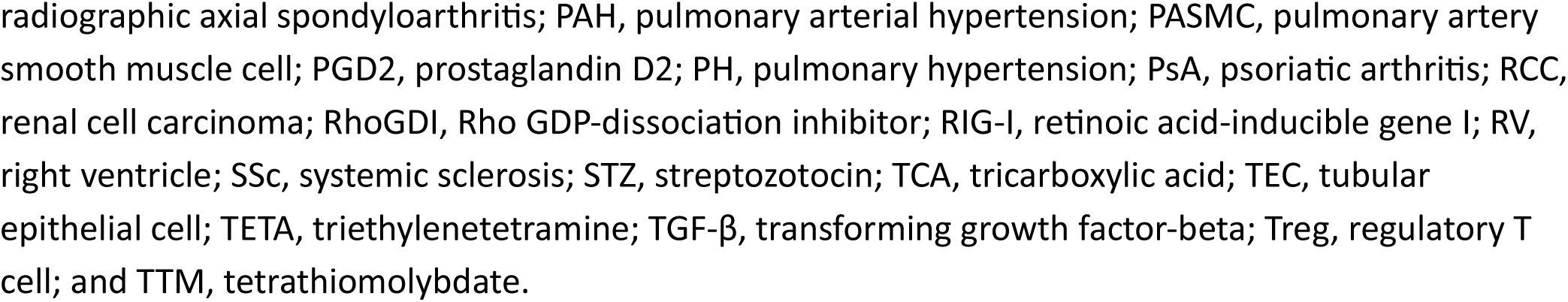
Candidate Therapeutic Interventions Targeting Key Pathways in PAH-Associated Kidney Dysfunction.

## Limitations

We acknowledge several key limitations in our study. First, our study analyzed autopsy-derived kidneys from a small cohort (5 control, 4 PAH), limiting statistical power and inference. Although autopsy tissue enables analysis of end-stage human disease, it represents a single time point and cannot establish temporal relationships; thus, our findings are purely descriptive and do not define causality. Second, the significant reduction in collecting duct populations in PAH kidneys limited nuclei available for differential expression analysis, precluding detailed transcriptional characterization of these segments. In chronic kidney disease, intercalated-to-principal cell transdifferentiation is proposed as a mechanism of collecting duct loss^47,48^. To assess whether this occurs in PAH nephropathy, we performed pseudotime trajectory and Notch signaling analyses, neither of which supported transdifferentiation (**Supplemental Figure 11**). However, definitive evaluation would require lineage-tracing studies. Third, our phosphoproteomic analyses identified reduced regulatory phosphosites in GSTA2, IDH1, and HAO2, but the functional consequences of these alterations remain to be experimentally validated. The HAO2–PEX5 models predict an altered targeting interface but do not establish changes in peroxisomal import or enzyme activity. Finally, we evaluated post-translational regulation only at the level of phosphorylation, and it is likely that other post-translational regulatory mechanisms also contribute to metabolic dysregulation in PAH nephropathy.

## Acknowledgements

We thank the University of Minnesota Genomics Center for performing the sequencing for the snRNA-seq experiment.

## Supplemental Materials

Expanded Methods

Supplemental Figures 1-11

Supplemental Table 1-3

## Non-Standard Abbreviations and Acronyms

AF3: AlphaFold 3
AKT1: AKT serine/threonine kinase 1
BCAA: branched-chain amino acid
CK2α: casein kinase 2 alpha
CKM: cardiovascular-kidney-metabolic
CSNK2A1: casein kinase 2 alpha 1
DBA: Dolichos biflorus agglutinin
DEG: differentially expressed gene
FDR: false-discovery rate
GSH: glutathione
GSK3A/B: glycogen synthase kinase 3 alpha/beta
GSTA2: glutathione S-transferase alpha 2
HAO2: 2-hydroxyacid oxidase 2
HIF-1: hypoxia-inducible factor 1
HLA: human leukocyte antigen
HPF: high-power field
IDH1: isocitrate dehydrogenase 1
IDO1: indoleamine 2,3-dioxygenase 1
IL: interleukin
KEA3: Kinase Enrichment Analysis 3
KEGG: Kyoto Encyclopedia of Genes and Genomes
LTL: Lotus tetragonolobus lectin
MAPK1/8: mitogen-activated protein kinase 1/8
MHC-I: major histocompatibility complex class I
NK: natural killer
OXPHOS: oxidative phosphorylation
PAH: pulmonary arterial hypertension
PCA: principal component analysis
PCT: proximal convoluted tubule
PEX5: peroxisomal biogenesis factor 5
pLDDT: predicted local distance difference test
PT: proximal tubule
PTS1: peroxisomal targeting signal 1
RAF1: Raf-1 proto-oncogene serine/threonine kinase
RMSD: root-mean-square deviation
RV: right ventricular
SASA: solvent-accessible surface area
snRNA-seq: single-nucleus RNA sequencing
SRPK3: SRSF protein kinase 3
STK3/4: serine/threonine-protein kinase ¾
TAL: thick ascending limb
TCA: tricarboxylic acid
TMT: tandem mass tag
UMAP: uniform manifold approximation and projection
VEGFA/VEGFR2: vascular endothelial growth factor A/receptor 2

