## Supplemental Data for "Multi-Omic Profiling Defines the Renal Mechanisms of Cardiovascular-Kidney-Metabolic Syndrome in Pulmonary Arterial Hypertension"

**Supplemental Figure 1. Single-nucleus RNA-sequencing quality-control metrics in control and PAH kidneys.** Violin plots show total RNA counts per nucleus (nCount_RNA), detected genes per nucleus (nFeature_RNA), and the percentages of mitochondrial (percent.mt) and ribosomal (percent.rb) transcripts after quality-control filtering. Horizontal bars and values above the distributions indicate group medians. PAH, pulmonary arterial hypertension.

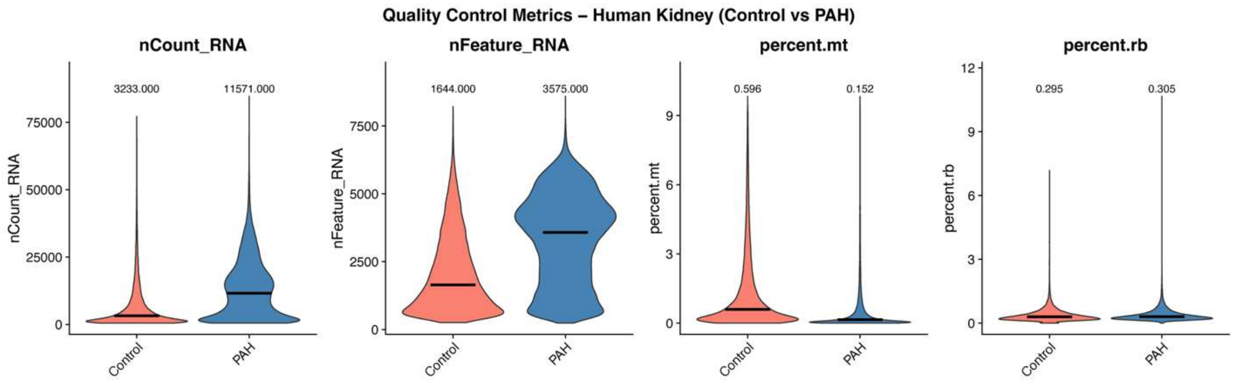

**Supplemental Figure 2. Single-nucleus RNA sequencing reveals altered cellular composition and extensive cell-type-specific transcriptional remodeling in PAH kidneys.** (A) Uniform manifold approximation and projection (UMAP) plots of 32,331 control nuclei from 5 patients and 56,993 PAH nuclei from 4 patients, representing 11 distinct renal cell types. Group-specific relative abundance values for each cell type are shown below each respective UMAP. (B) Heatmap showing representative marker-gene expression used to annotate renal cell types. (C) Relative abundance of each cell type by patient in control and PAH kidneys. Each point represents one patient; horizontal lines indicate group medians. P values shown were calculated using two-sided Mann–Whitney U tests. (D) Principal component analysis of sample-level transcriptomes, colored by disease group; shaded ellipses indicate group dispersion. (E) Cell-type-resolved differential expression in PAH versus control kidneys. Red and blue indicate genes increased and decreased in PAH, respectively; points denote differentially expressed genes meeting adjusted $P < 0.05$ and absolute log2 fold change > 0.5. Differential expression was performed on sample-pseudobulked counts using DESeq2, with Benjamini–Hochberg correction. DCT, distal convoluted tubule; EC, endothelial cell; FB, fibroblast; IC, intercalated cell; LC, lymphocyte; MP, macrophage; PC, principal cell; POD, podocyte; PT, proximal tubule; PTL, parietal cell; TAL, thick ascending limb.

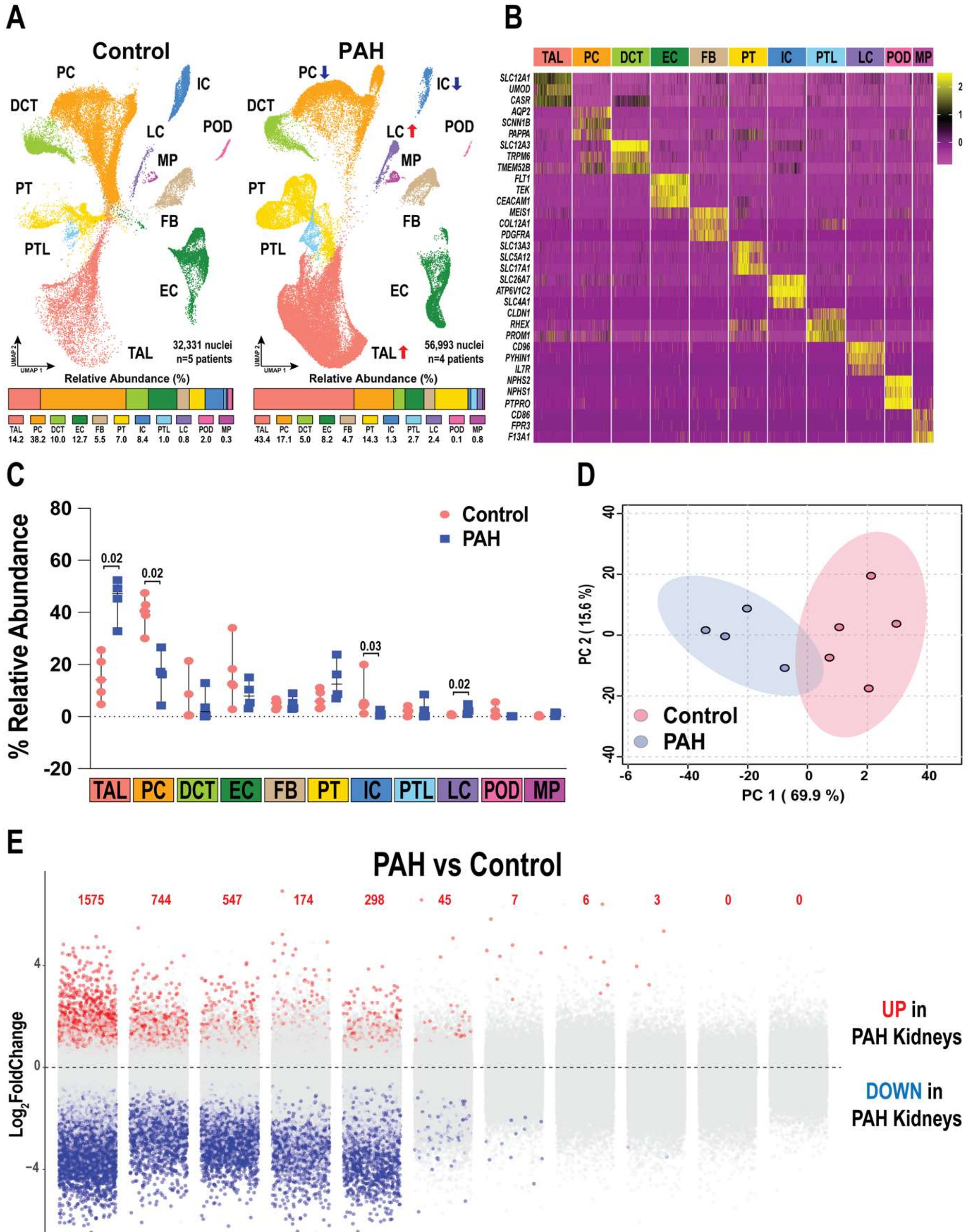
A
Control
PAH
PC
IC
DCT
LC
POD
MP
PT
FB
PTL
EC
TAL
32,331 nuclei
n=5 patients
56,993 nuclei
n=4 patients
Relative Abundance (%)
TAL 14.2 PC 38.2 DCT 10.0 EC 12.7 FB 5.5 PT 7.0 IC 8.4 PTL 1.0 LC 0.8 POD 2.0 MP 0.3
TAL 43.4 PC 17.1 DCT 5.0 EC 8.2 FB 4.7 PT 14.3 IC 1.3 PTL 2.7 LC 2.4 POD 0.1 MP 0.8
B
TAL PC DCT EC FB PT IC PTL LC POD MP
SLC12A1 UMOD CASR AQP2 SCNN1B PAPPA SLC12A3 TRPM6 TMEM52B FLT1 TEK CEACAM1 MEIS1 COL12A1 PDGFRA SLC13A3 SLC5A12 SLC17A1 SLC26A7 ATP6V1C2 SLC4A1 CLDN1 RHEX PROM1 CD96 PYHIN1 IL7R NPHS2 NPHS1 PTPRO CD86 FPR3 F13A1
C
% Relative Abundance
Control
PAH
0.02
0.02
0.03
0.02
D
PC 2 ( 15.6 %)
PC 1 ( 69.9 %)
Control
PAH
E
PAH vs Control
1575 744 547 174 298 45 7 6 3 0 0
Log2FoldChange
UP in PAH Kidneys
DOWN in PAH Kidneys

**Supplemental Figure 3. Pathway enrichment among genes downregulated in proximal tubule and thick ascending limb cells from PAH kidneys.** (A) Top Kyoto Encyclopedia of Genes and Genomes (KEGG; A) and (B) WikiPathways terms enriched among genes downregulated in PAH proximal tubule cells relative to controls. (C) Top KEGG (C) and (D) WikiPathways terms enriched among genes downregulated in PAH thick ascending limb cells relative to controls. Dot size indicates the number of genes, color represents −log10(false-discovery rate [FDR]), and the x-axis shows fold enrichment.

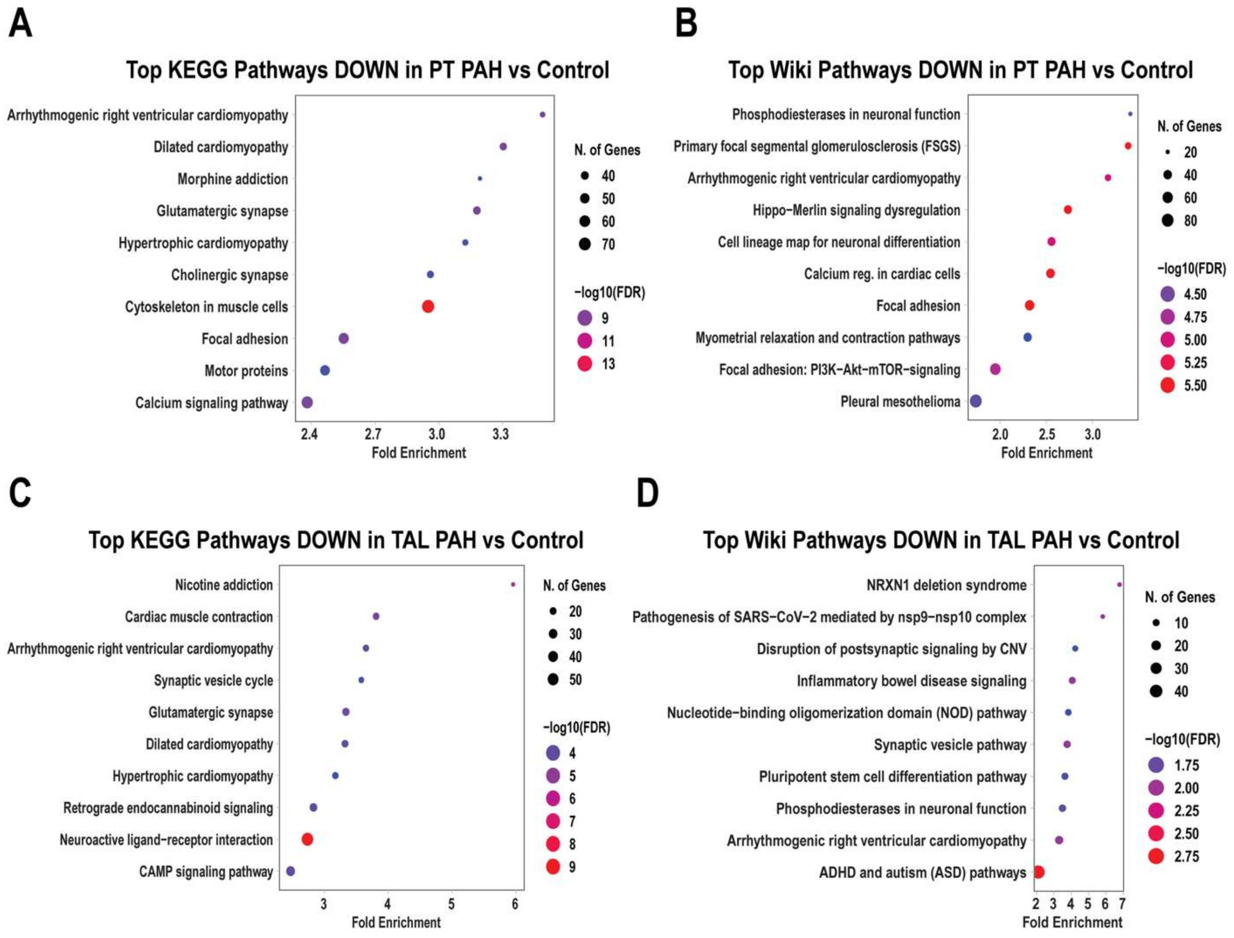

**Supplemental Figure 4. Pathway enrichment among genes downregulated in PAH lymphocytes.** (A) Top KEGG and (B) WikiPathways terms enriched among genes downregulated in PAH lymphocytes relative to controls. Dot size indicates the number of genes, color represents −log10(false-discovery rate [FDR]), and the x-axis shows fold enrichment.

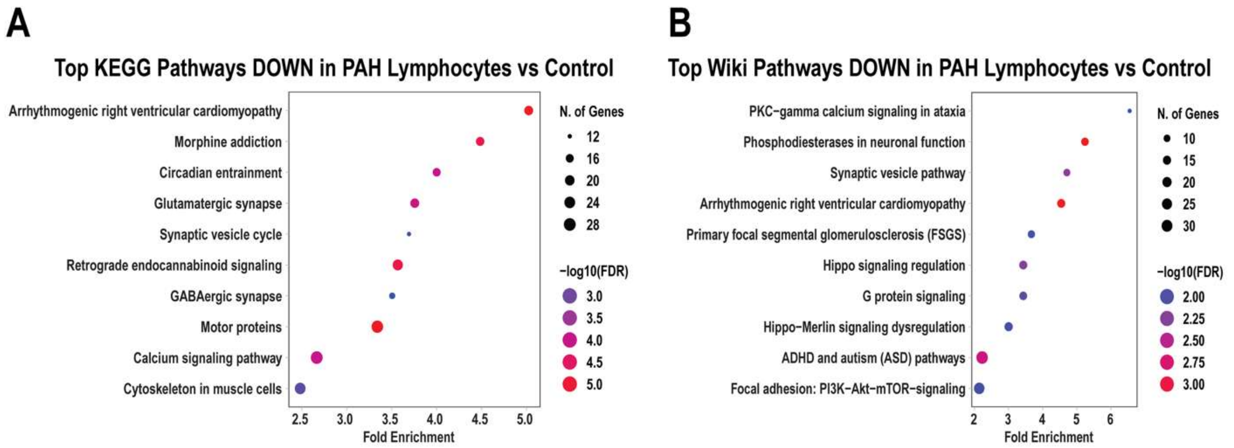

**Supplemental Figure 5. Lymphocyte subtypes in control and PAH kidneys.** Uniform manifold approximation and projection (UMAP) plots show lymphocyte nuclei from control and PAH kidneys, colored by annotated subtype: CD4+ T cells, CD8+ T cells, natural killer (NK) cells, and B cells. Each point represents one nucleus.

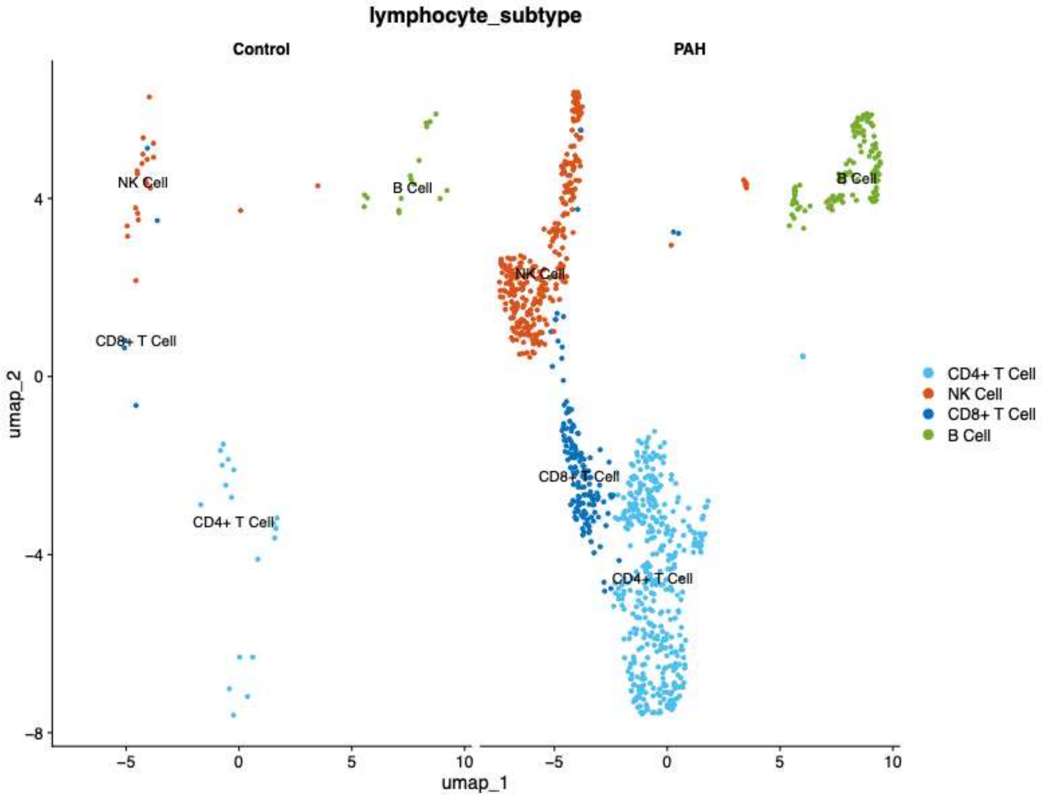

**Supplemental Figure 6. Mitochondrial proteomic pathway remodeling in PAH kidneys.** (A) WikiPathways terms enriched among proteins decreased or (B) increased in mitochondria-enriched fractions from PAH kidneys relative to controls. Dot size indicates the number of pathway-associated proteins, color represents −log10(false-discovery rate [FDR]), and the x-axis shows fold enrichment.

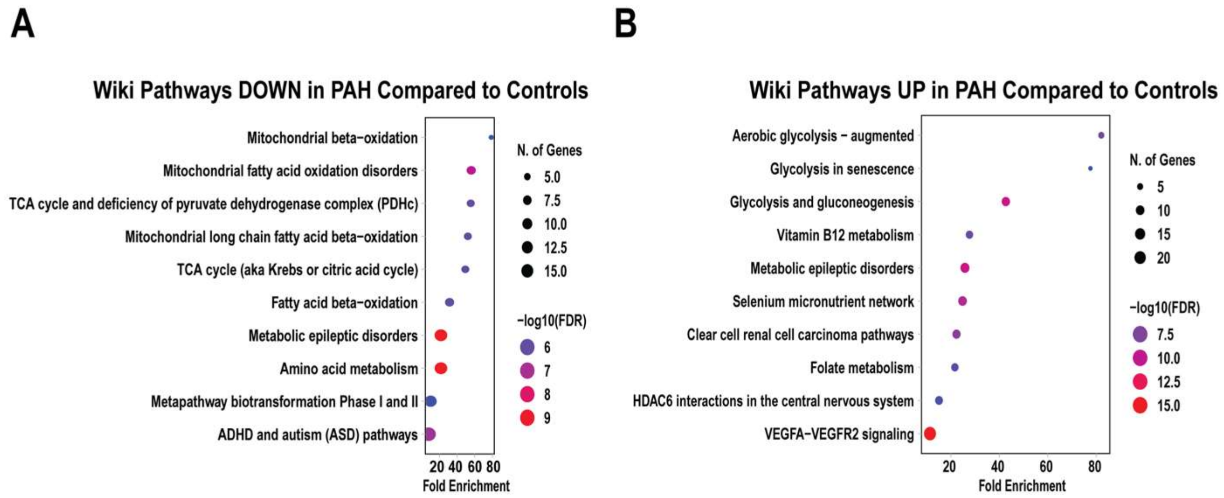

**Supplemental Figure 7. Cytoplasmic proteomic pathways increased in PAH kidneys.** (A) Top KEGG and (B) WikiPathways terms enriched among proteins increased in cytoplasmic fractions from PAH kidneys relative to controls. Dot size indicates the number of pathway-associated proteins, color represents −log10(false-discovery rate [FDR]), and the x-axis shows fold enrichment.

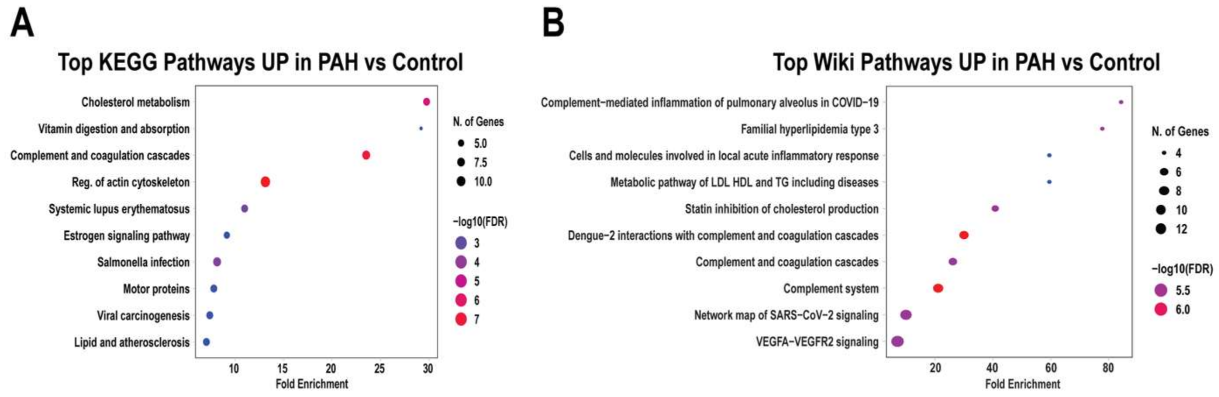

**Supplemental Figure 8. PAH kidneys exhibit a distinct phosphoproteomic signature, predicted kinase dysregulation, and phosphorylation-associated remodeling of metabolic enzymes.** (A) Unsupervised hierarchical clustering of the 500 most variable phosphoproteins in control and PAH kidneys. (B) Principal component analysis of phosphoproteomic profiles; points represent patients, colors indicate disease group, and shaded ellipses indicate group dispersion. (C) Kinase Enrichment Analysis 3 (KEA3) MeanRank predictions for kinases associated with phosphoproteins increased or (D) decreased in PAH versus controls. Bar segments indicate the contributing kinase–substrate databases. (E) AlphaFold 3 structural predictions of native (tan) and phosphorylated (blue) 2-hydroxyacid oxidase 2 (HAO2; pS349) in complex with the peroxisomal import receptor PEX5 (salmon). Overviews identify the PEX5 binding pocket, and enlarged views show the predicted intermolecular hydrogen-bond networks formed by the HAO2 C-terminal PTS1 motif in the native and pS349 models. Asterisks identify PEX5 residues with phosphorylation-state-specific hydrogen-bond contacts. (F) AlphaFold 3 structural predictions comparing native (tan) and phosphorylated (blue) glutathione S-transferase alpha 2 (GSTA2; pS37). Overlays show the phosphosite and catalytic Tyr9, surface maps show phosphorylation-associated changes in electrostatic potential, and the enlarged view shows distances from S37/pS37 to the glutathione-binding site and Tyr9. (G) AlphaFold 3 structural predictions comparing the native (tan) and phosphorylated (blue) isocitrate dehydrogenase 1 homodimer (IDH1; pS94). Overlays show pS94 and catalytic Arg100 and Arg132, surface maps show phosphorylation-associated changes in solvent-accessible surface area (SASA), and the enlarged view shows distances from S94/pS94 to Arg100 and Arg132. Electrostatic potentials were calculated over a fixed −10 to +10 kcal/mol·e range. GSK3A/B, glycogen synthase kinase 3α/β; CSNK2A1, casein kinase 2 alpha 1; MAPK1/8, mitogen-activated protein kinase 1/8; RAF1, Raf-1 proto-oncogene serine/threonine kinase; SRPK3, SRSF protein kinase 3; STK3/4, serine/threonine-protein kinase 3/4.

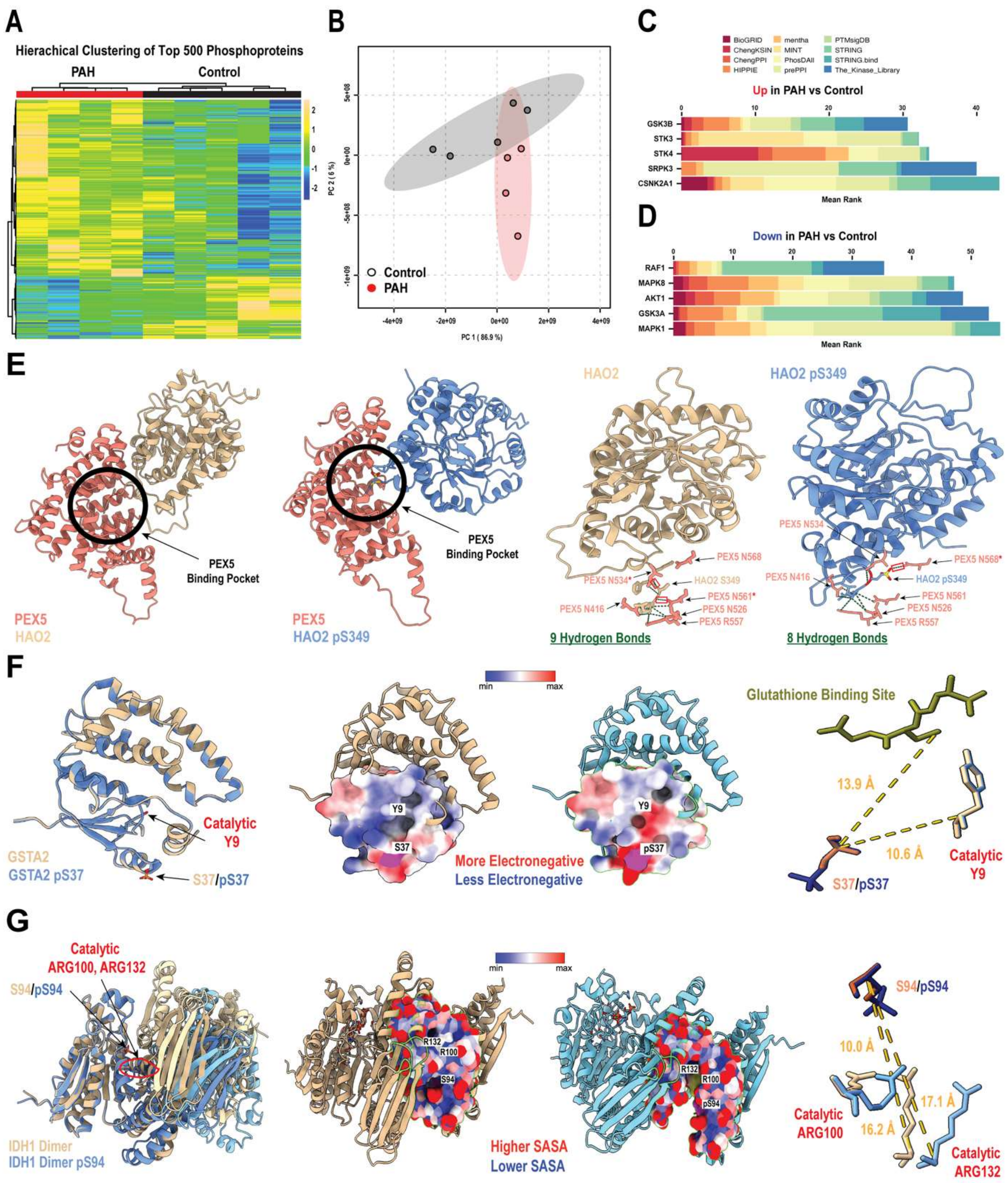
A
Hierachical Clustering of Top 500 Phosphoproteins
PAH
Control
B
PC 2 ( 6 %)
PC 1 ( 86.9 %)
Control
PAH
C
BioGRID
ChengKSIN
ChengPPI
HIPPIE
mentha
MINT
PhosDAll
prePPI
PTMsigDB
STRING
STRING.bind
The_Kinase_Library
Up in PAH vs Control
GSK3B
STK3
STK4
SRPK3
CSNK2A1
Mean Rank
D
Down in PAH vs Control
RAF1
MAPK8
AKT1
GSK3A
MAPK1
Mean Rank
E
PEX5 Binding Pocket
PEX5
HAO2
PEX5
HAO2 pS349
HAO2
HAO2 pS349
PEX5 N568
PEX5 N534*
HAO2 S349
PEX5 N416
PEX5 N561*
PEX5 N526
PEX5 R557
9 Hydrogen Bonds
PEX5 N534
PEX5 N568*
PEX5 N416
HAO2 pS349
PEX5 N561
PEX5 N526
PEX5 R557
8 Hydrogen Bonds
F
Catalytic Y9
GSTA2
GSTA2 pS37
S37/pS37
min
max
Y9
S37
pS37
More Electronegative
Less Electronegative
Glutathione Binding Site
13.9 Å
10.6 Å
S37/pS37
Catalytic Y9
G
Catalytic ARG100, ARG132
S94/pS94
IDH1 Dimer
IDH1 Dimer pS94
min
max
R132
R100
S94
pS94
Higher SASA
Lower SASA
10.0 Å
17.1 Å
16.2 Å
Catalytic ARG100
Catalytic ARG132

**Supplemental Figure 9. Kynurenine pathway protein abundance in control and PAH kidneys.** Combined cytoplasmic and mitochondrial protein abundance is shown for AADAT, ACMSD, AFMID, HAAO, KMO, and KYNU. Boxes show medians and interquartile ranges, and each point represents one patient. Exact P values were calculated using two-sided Mann–Whitney tests. AADAT, aminoadipate aminotransferase; ACMSD, aminocarboxymuconate-semialdehyde decarboxylase; AFMID, arylformamidase; HAAO, 3-hydroxyanthranilate 3,4-dioxygenase; KMO, kynurenine 3-monooxygenase; KYNU, kynureninase.

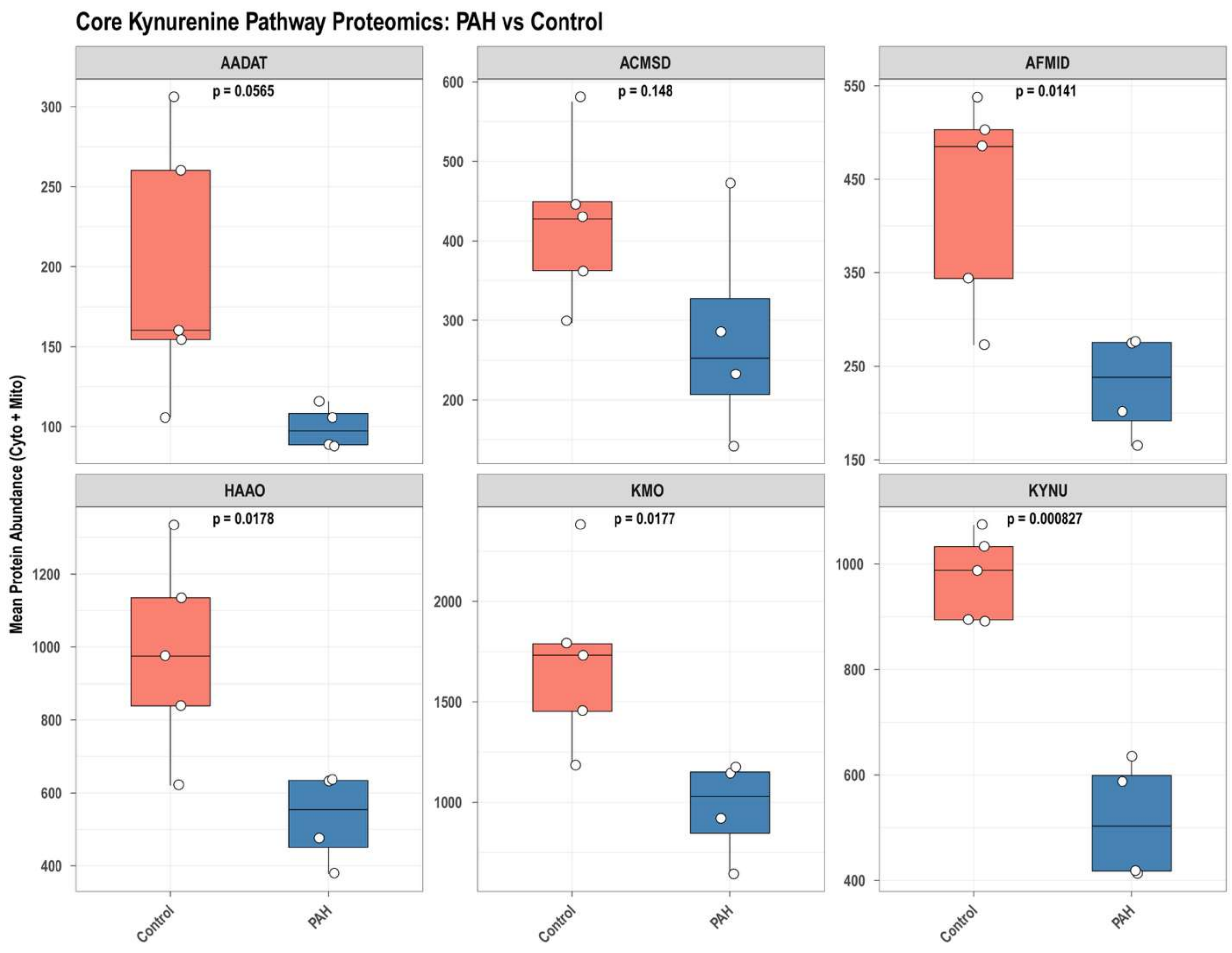

**Supplemental Figure 10. Endothelial-cell and fibroblast pathway remodeling in PAH kidneys.** (A) Top KEGG and WikiPathways terms enriched among genes upregulated in PAH endothelial cells. (B) Pseudobulked fatty acid metabolism and cuproptosis module scores in control and PAH endothelial cells. (C) Top KEGG and WikiPathways terms enriched among genes upregulated in PAH fibroblasts. (D) Pseudobulked glycolysis and ferroptosis module scores in control and PAH fibroblasts. In enrichment plots, dot size indicates the number of genes, color represents −log10(false-discovery rate [FDR]), and the x-axis shows fold enrichment. In module-score plots, each point represents one patient and horizontal bars indicate medians; exact P values were calculated using two-sided Mann–Whitney tests.

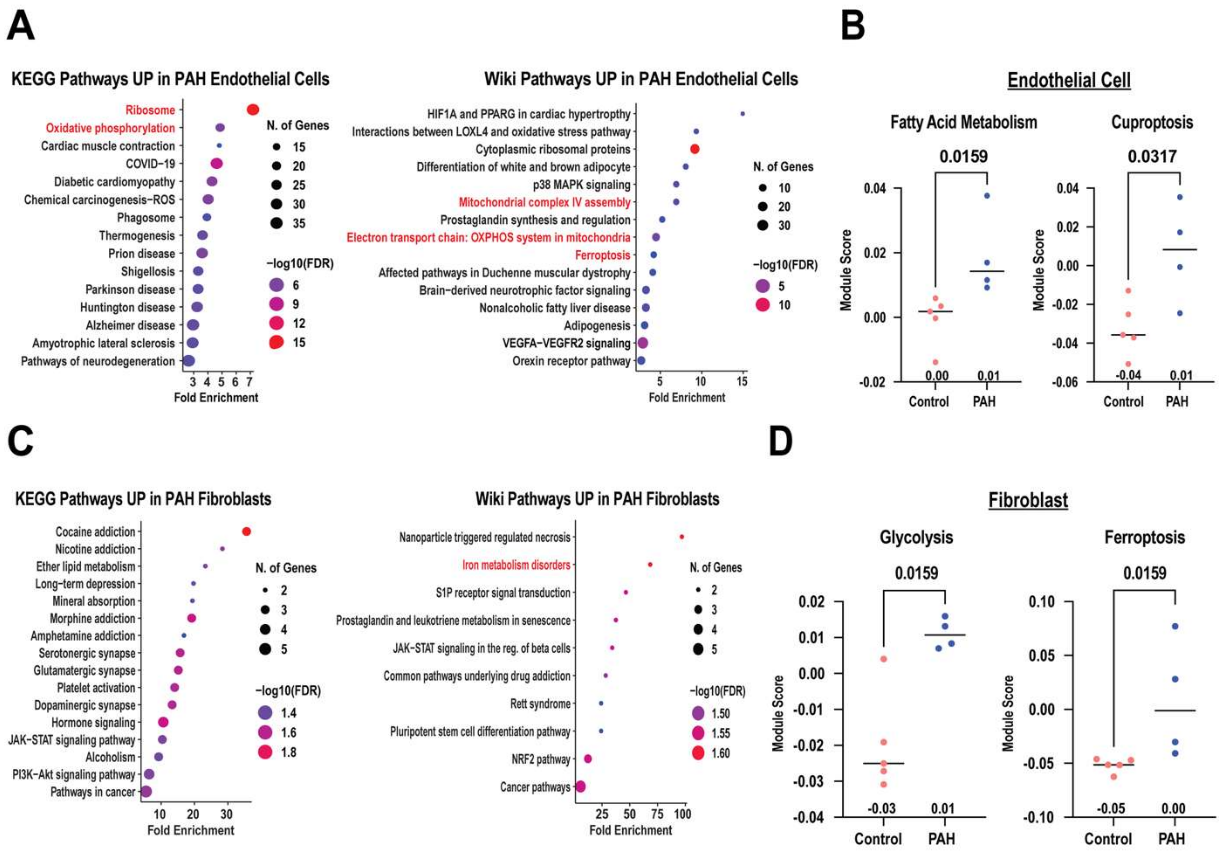

**Supplemental Figure 11. Principal-cell to intercalated-cell trajectory analysis in control and PAH kidneys.** (A) Uniform manifold approximation and projection (UMAP) plots show principal-cell (PC) and intercalated-cell (IC) nuclei by disease group; labels indicate the number of nuclei in each population. (B) Nuclei are colored by Monocle 3 pseudotime. (C) Nuclei are colored by the Notch/PC module score. (D) Patient-level mean IC pseudotime and mean Notch/PC module score in ICs. Point size indicates the number of ICs per sample, and horizontal bars indicate medians. Exact P values were calculated using two-sided Mann–Whitney tests.

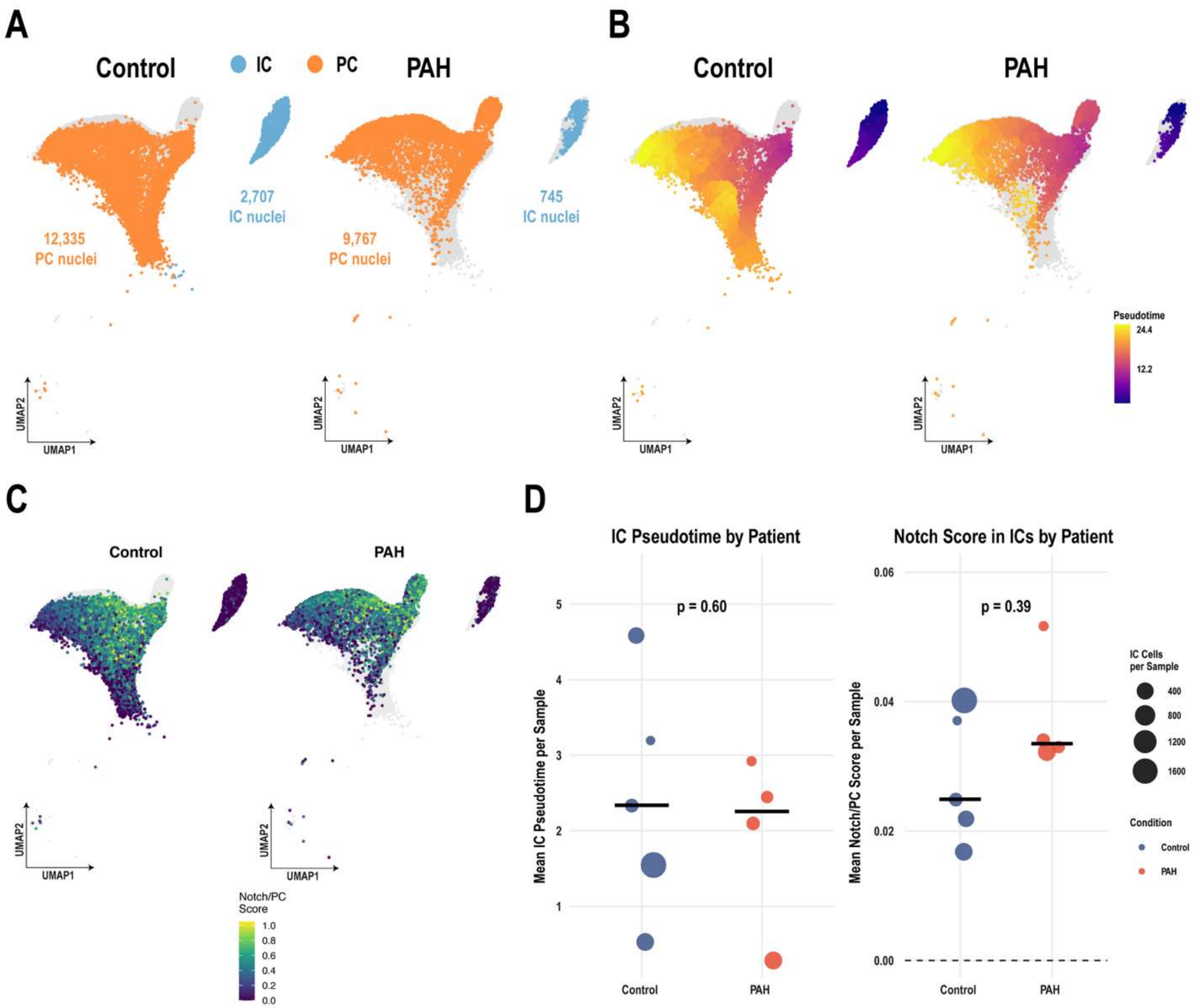

### Supplemental Table 1A: Clinical Characteristics of Patient Cohort

| | | | | | | | ECHOCARDIOGRAPHY # | | | | | RIGHT HEART CATHETERIZATION ## | | | | | | NT-proBNP | |
|---|---|---|---|---|---|---|---|---|---|---|---|---|---|---|---|---|---|---|---|
| **Category** | **Diagnosis** | **Age (yr)** | **Sex** | **WHO Class** | **Height (cm)** | **Weight (kg)** | **Interval, † (days)** | **LVEF (%)** | **RVSP (mmHg)** | **TAPSE (mm)** | **CI (L/min/$m^2$)** | **Interval, ‡ (days)** | **mPAP (mmHg)** | **RAP (mmHg)** | **CI (L/min/$m^2$)** | **CO (L/min)** | **PVR (dyne·s/$cm^5$)** | **Interval, § (days)** | **NT-proBNP (pg/mL)** |
| CTRL1 | Sudden death (CAD)* | 52 | F | N/A | 175 | 72 | N/A | N/A | N/A | N/A | N/A | N/A | N/A | N/A | N/A | N/A | N/A | N/A | N/A |
| CTRL2 | Sudden death (CAD)* | 29 | M | N/A | 160 | 67.8 | N/A | N/A | N/A | N/A | N/A | N/A | N/A | N/A | N/A | N/A | N/A | N/A | N/A |
| CTRL3 | Sudden death (CAD)* | 65 | M | N/A | N/A | N/A | N/A | N/A | N/A | N/A | N/A | N/A | N/A | N/A | N/A | N/A | N/A | N/A | N/A |
| CTRL4 | Sudden death* | 48 | F | N/A | N/A | N/A | N/A | N/A | N/A | N/A | N/A | N/A | N/A | N/A | N/A | N/A | N/A | N/A | N/A |
| CTRL5 | Sudden death (drug poisoning)* | 49 | F | N/A | N/A | N/A | N/A | N/A | N/A | N/A | N/A | N/A | N/A | N/A | N/A | N/A | N/A | N/A | N/A |
| PAH1 | HPAH | 61 | F | III | 163 | 43 | 7 | 45 | 100 | 16 | 1.73 | 2,085 | 66 | 7 | 2.90 | 4.5 | 1,012 | 11 | 8,386 |
| PAH2 | SSc-PAH | 68 | F | IV | 163 | 58.5 | 761 | 45 | N/A | 13 | 1.78 | 263 | 43 | 6 | 1.82 | 3.07 | 1,042 | 4,574 | 214 |
| PAH3 | IPAH | 82 | M | IV | 173 | 88 | 229 | 60 | 27 | 18 | 2.01 | 267 | 25 | 2 | 2.57 | 5.2 | 308 | 5 | 3,167 |
| PAH4 | HPAH | 39 | F | IV | 157 | 64 | 6 | 60 | N/A | 13 | 1.40 | N/A | N/A | N/A | N/A | N/A | N/A | 6 | 904 |

**Supplemental Table 1B.** Medications and laboratory values for kidney cohort subjects.

| | VASOACTIVE THERAPY / DIURETICS | | | | | COMPLETE BLOOD COUNT (×10⁹/L) | | | | | | | RENAL FUNCTION | | |
|---|---|---|---|---|---|---|---|---|---|---|---|---|---|---|---|
| Category | ERA (duration) | PDE5 Inhibitor | Epopro-stenol | Selex-ipag | Lasix (duration) | Interval,† (days) | WBC | Neut. | Lymph. | Mono. | Eos. | Baso. | Interval,‡ (days) | Creatinine (µM) | eGFR (mL/min/1.73m²) |
| CTRL1 | N/A | N/A | N/A | N/A | N/A | N/A | N/A | N/A | N/A | N/A | N/A | N/A | N/A | N/A | N/A |
| CTRL2 | N/A | N/A | N/A | N/A | N/A | N/A | N/A | N/A | N/A | N/A | N/A | N/A | N/A | N/A | N/A |
| CTRL3 | N/A | N/A | N/A | N/A | N/A | N/A | N/A | N/A | N/A | N/A | N/A | N/A | N/A | N/A | N/A |
| CTRL4 | N/A | N/A | N/A | N/A | N/A | N/A | N/A | N/A | N/A | N/A | N/A | N/A | N/A | N/A | N/A |
| CTRL5 | N/A | N/A | N/A | N/A | N/A | N/A | N/A | N/A | N/A | N/A | N/A | N/A | N/A | N/A | N/A |
| PAH1 | Yes (>2 yr) | Yes (>2 yr) | No | Yes (>2 yr) | Yes (>2 yr) | 1 | 16.9 | 14.7 | 1.2 | 0.9 | 0.1 | 0.0 | 1 | 103 | 51 |
| PAH2 | Yes (>2 yr, 1 mo) | Yes (>2 yr, 3 mo) | No | No | Yes (9 mo) | 17 | 5.8 | 4.1 | 1.2 | 0.5 | 0.0 | 0.0 | 17 | 124 | 39 |
| PAH3 | Yes (>2 yr, 3 mo) | Yes (>2 yr, 6 mo) | No | No | Yes (>2 yr) | 3 | 8.0 | 4.2 | 2.7 | 0.8 | 0.2 | 0.0 | 3 | 139 | 40 |
| PAH4 | Yes (2 days) | Yes (10 days) | Yes (10 days) | No | Yes (9 days) | 1 | 9.0 | 6.5 | 1.7 | 0.6 | 0.1 | 0.0 | 1 | 53 | 115 |

†Interval between complete blood count and tissue sampling.

‡Interval between creatinine measurement and tissue sampling.

*Early autopsy performed following sudden death. None of these subjects had any past medical history. No macroscopic signs of chronic right or left heart dysfunction were observed; all subjects exhibited normal heart size, no cardiac hypertrophy (left and right ventricular wall diameter 1.3–1.5 cm and 0.2–0.5 cm, respectively), and normal pulmonary arteries.

Abbreviations: CAD, coronary artery disease; CI, cardiac index; CO, cardiac output; eGFR, estimated glomerular filtration rate; ERA, endothelin receptor antagonist; Eos., eosinophils; HPAH, heritable pulmonary arterial hypertension; IPAH, idiopathic pulmonary arterial hypertension; LVEF, left ventricular ejection fraction; Lymph., lymphocytes; mPAP, mean pulmonary artery pressure; Mono., monocytes; N/A, not applicable or not available; Neut., neutrophils; NT-proBNP, N-terminal prohormone of brain natriuretic peptide; PDE5i, phosphodiesterase-5 inhibitor; PVR, pulmonary vascular resistance; RAP, right atrial pressure; RVSP, right ventricular systolic pressure; SSc-PAH, systemic sclerosis–associated pulmonary arterial hypertension; TAPSE, tricuspid annular plane systolic excursion; WBC, white blood cell count; WHO, World Health Organization.

**Supplemental Table 2. Single-nucleus RNA sequencing library quality metrics.**

| Sample | Nuclei Sequenced, n | Median Genes | Reads Per Nucleus, n | Total Reads | Mapped to Genome, % |
|---|---|---|---|---|---|
| CTRL42 | 13,582 | 668 | 72,260 | 981,436,566 | 92.6 |
| CTRL43 | 5,352 | 678 | 176,421 | 944,203,496 | 93.1 |
| CTRL45 | 11,091 | 1,655 | 83,636 | 927,602,971 | 92.6 |
| CTRL51 | 15,476 | 2,308 | 56,942 | 881,240,173 | 93.7 |
| CTRL55 | 9,350 | 562 | 94,176 | 885,590,893 | 93.2 |
| PAH119 | 22,931 | 4,059 | 37,372 | 856,973,951 | 94.5 |
| PAH280 | 15,228 | 3,670 | 55,636 | 847,218,001 | 94.1 |
| PAH305 | 13,031 | 2,060 | 61,363 | 799,618,158 | 93.5 |
| PAH306 | 16,755 | 4,002 | 52,024 | 871,659,978 | 94.3 |

**Supplemental Table 3. AlphaFold3 Structural Predictions for Differentially Phosphorylated Proteins in PAH Kidneys**

Structural metrics for HAO2 (pS349), GSTA2 (pS37), and IDH1 (pS94) comparing native and phosphorylated AlphaFold 3 predictions. HAO2 was modeled as its C-terminal peroxisomal targeting signal 1 (PTS1) peptide bound to PEX5; GSTA2 and IDH1 were modeled with GSH and $NADP^+$, respectively.

| | **HAO2 (pS349)** | | | **GSTA2 (pS37)** | | | **IDH1 (pS94)** | | |
|---|---|---|---|---|---|---|---|---|---|
| **Measurement** | **Native** | **Phospho** | **Δ** | **Native** | **Phospho** | **Δ** | **Native** | **Phospho** | **Δ** |
| ***Enzyme Function*** | | | | | | | | | |
| Classification | FAD-dependent oxidoreductase | | | GSH-conjugating transferase | | | $NADP^+$-dependent oxidoreductase | | |
| Physiological ligand / binding partner | PEX5 | | | GSH | | | $NADP^+$ | | |
| *Phosphoproteomics* | | | | | | | | | |
| PAH/Control ratio | | | 0.260 | | | 0.323 | | | 0.313 |
| Adjusted P-value | | | 0.036 | | | 0.096 | | | 0.072 |
| *Structural Quality* | | | | | | | | | |
| Structural RMSD (Å) | — | 0.631 (PTS1 backbone) | | — | 0.116 | | — | 0.520 | |
| pLDDT at phosphosite[†] | 97.63 | 91.36 | −6.27 | 98.56 | n/a[†] | — | 93.22 | 93.19 | −0.03 |
| pLDDT range (modeled complex) | 26.9–99.0 | 29.8–99.0 | | 44.1–99 | 38–99 | | | | |
| *Key Site and Interface Metrics* | | | | | | | | | |
| HAO2 PTS1 RMSD, residues 349–351 (Å) | — | 0.324 | | | | | | | |

| | | | | | | | | | |
|---|---|---|---|---|---|---|---|---|---|
| S349 Cα displacement (Å) | — | 0.461 | | | | | | | |
| HAO2–PEX5 buried interface area (Å²) | 584.06 | 602.32 | +18.26 (+3.1%) | | | | | | |
| HAO2–PEX5 H-bonds | 13 | 14 | +1 | | | | | | |
| Steric clashes at interface | 1 | 3 | +2 | | | | | | |
| TYR9 (G-site catalytic) | | | | 98.88 | 98.61 | −0.27 | | | |
| ARG100 (isocitrate binding) | | | | | | | 86.52 | 91.56 | +5.04 |
| ARG132 (catalytic) | | | | | | | 89.61 | 94.81 | +5.20 |
| *Local Electrostatics, 10 Å Zone (kcal/mol·e)* | | | | | | | | | |
| Mean potential | | | | +0.29 | −0.89 | −1.18 | −0.77 | −2.06 | −1.29 |
| Minimum potential | | | | −8.43 | −19.77 | −11.34 | −14.84 | −17.75 | −2.91 |
| Maximum potential | | | | +17.03 | +8.74 | −8.29 | +14.76 | +8.30 | −6.46 |
| *Whole-Surface Electrostatics (kcal/mol·e)* | | | | | | | | | |
| Mean potential | | | | −0.43 | −0.96 | −0.53 | −1.09‡ | −1.53‡ | −0.44‡ |
| Chain B mean potential§ | | | | | | | −1.10 | −1.09 | +0.01 |
| *Solvent-Accessible Surface Area (Å²)* | | | | | | | | | |
| Local SASA, 10 Å zone | 1,425.2 | 1,636.0 | +14.8% | 4,730 | 4,784 | +1.1% | 6,441 | 7,336 | +13.9% |
| Global SASA | 15,492 | 15,534 | +0.27% | 12,159 | 12,155 | −0.03% | 20,568 | 20,616 | +0.23% |
| Hydrophobic SASA | | | | 10,228 | 10,152 | −0.74% | 17,618 | 17,791 | +0.98% |

| | | | | | | | | | |
|---|---|---|---|---|---|---|---|---|---|
| *Key Site Distances (Å)* | | | | | | | | | |
| Phosphosite → key residue/atom | 3.347 (PEX5 A533 Cβ) | 1.491 (PEX5 A533 Cβ) | −1.856 | 10.595 (Y9) | 10.584 (Y9) | −0.011 | 10.011 (R100) | 9.970 (R100) | −0.041 |
| S94 → R132 (IDH1 only) | | | | | | | 16.205 | 17.132 | +0.927 |
| *IDH1 Dimer Interface* | | | | | | | | | |
| Buried interface area (Å²) | | | | | | | 3,024.5 | 3,001.7 | −22.8 |
| Cross-interface H-bonds | | | | | | | 3,563 | 3,835 | +272 |
| Cross-interface contacts | | | | | | | 195,240 | 194,317 | −923 |

† Phosphosite pLDDT values were taken from Cα B_iso_or_equiv fields in representative AlphaFold 3 CIF files; n/a indicates that a value was not extractable for the modified residue.

‡ IDH1 whole-surface electrostatics reported for chain A (phosphorylated subunit).

§ Chain B is the unmodified partner subunit in the IDH1 homodimer; unchanged electrostatics confirm subunit-specific phosphorylation effects.

For GSTA2 and IDH1, SASA measurements use no-ligand models to avoid cofactor surface interference. HAO2 SASA and interface measurements use focused PEX5–HAO2 PTS1 models. Empty cells indicate that a measurement was not applicable or was not performed for that protein.

RMSD, root mean square deviation; pLDDT, predicted local distance difference test; SASA, solvent-accessible surface area. Δ = phospho − native.
